# The snoGloBe interaction predictor reveals a broad spectrum of C/D snoRNA RNA targets

**DOI:** 10.1101/2021.09.14.460265

**Authors:** Gabrielle Deschamps-Francoeur, Sonia Couture, Sherif Abou-Elela, Michelle S. Scott

## Abstract

Box C/D small nucleolar RNAs (snoRNAs) are a conserved class of RNA known for their role in guiding ribosomal RNA 2’-O-ribose methylation. Recently, C/D snoRNAs were also implicated in regulating the expression of non-ribosomal genes through different modes of binding. Large scale RNA-RNA interaction datasets detect many snoRNAs binding messenger RNA, but are limited by specific experimental conditions. To enable a more comprehensive study of C/D snoRNA interactions, we created snoGloBe, a human C/D snoRNA interaction predictor based on a gradient boosting classifier. SnoGloBe considers the target type, position and sequence of the interactions, enabling it to outperform existing predictors. Interestingly, for specific snoRNAs, snoGloBe identifies strong enrichment of interactions near gene expression regulatory elements including splice sites. Abundance and splicing of predicted targets were altered upon the knockdown of their associated snoRNA. Strikingly, the predicted snoRNA interactions often overlap with the binding sites of functionally related RNA binding proteins, reinforcing their role in gene expression regulation. SnoGloBe is also an excellent tool for discovering viral RNA targets, as shown by its capacity to identify snoRNAs targeting the heavily methylated SARS-CoV-2 RNA. Overall, snoGloBe is capable of identifying experimentally validated binding sites and predicting novel sites with shared regulatory function.

## INTRODUCTION

Small nucleolar RNAs (snoRNAs) are a conserved class of noncoding RNA required for rRNA modification, processing, and assembly (1). In addition, snoRNAs contribute to spliceosome biogenesis by guiding the modification of small nuclear RNA (snRNA) (2). To carry out these functions, deemed canonical, they assemble in ribonucleoprotein (snoRNP) complexes which provide them stability and catalytic activity. SnoRNAs are split in two families based on their structure, conserved motifs, interacting proteins and modification type. Box C/D snoRNAs guide 2’-O-ribose methylation while box H/ACA snoRNAs guide pseudouridylation of RNA (3, 4). SnoRNAs of both families identify their modification targets through base pairing between the snoRNA guide sequence and the sequence flanking the modification sites (5).

Box C/D snoRNAs usually range between 50-100 nucleotides in length (6) and are characterized by their conserved motifs: the boxes C/C’ (RUGAUGA) and D/D’ (CUGA) (Figure 1A). They interact with core binding proteins SNU13, NOP56, NOP58 and the methyltransferase fibrillarin (FBL) to form the C/D snoRNP. Box C/D snoRNAs guide their catalytic partner to the modification site using sequence complementarity to the region upstream of the boxes D and D’, called the antisense element (ASE) (3, 7). Box C/D snoRNA ASEs range between 10 and 20 nucleotides in length and are deemed essential for rRNA modification. However, many C/D snoRNAs have no identified canonical modification target and are referred to as orphan snoRNAs. Interestingly, some snoRNAs were also described as guiding the modification of messenger RNAs (mRNAs) expanding the confines of potential RNA targets (8).

**Figure 1.**
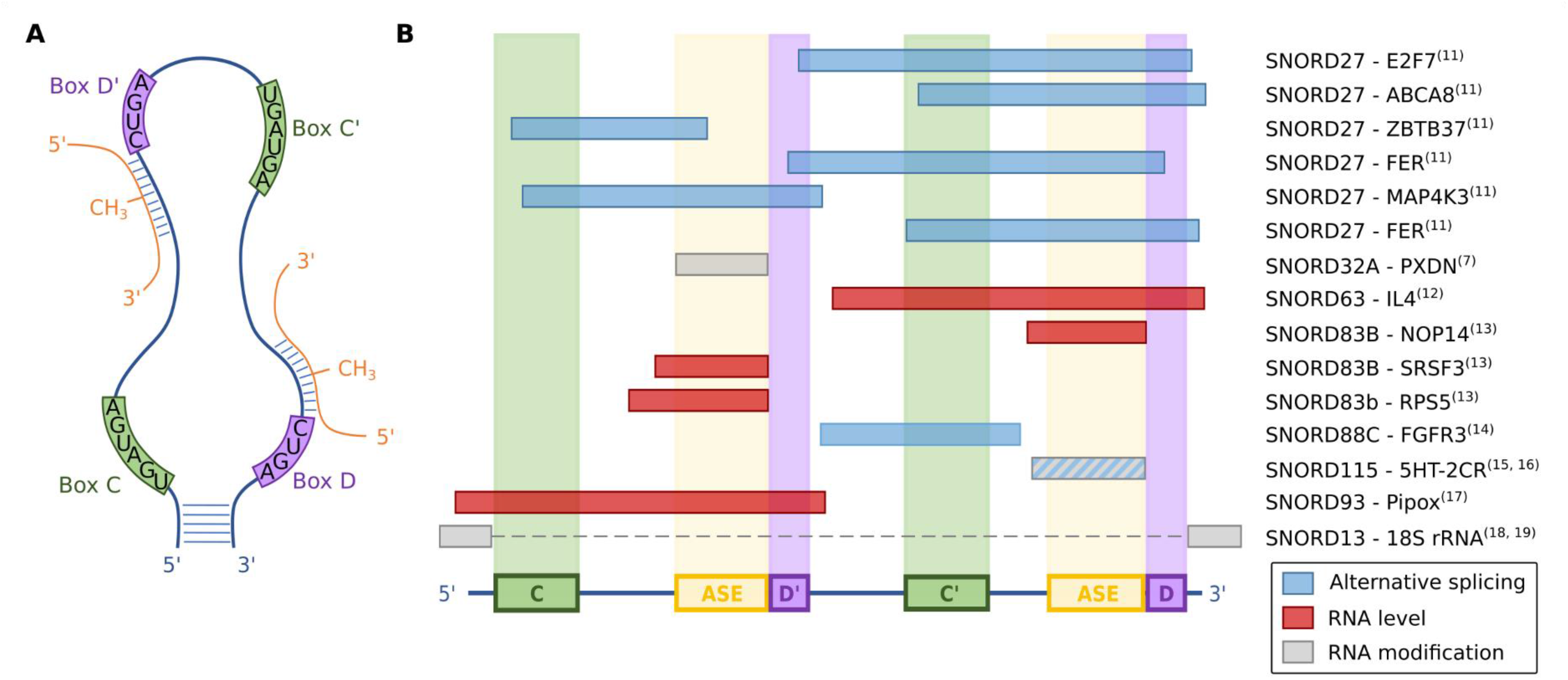
Box C/D snoRNA characteristics and interactions. A) Box C/D snoRNAs have well conserved patterns, called boxes C/C’ (green) and D/D’ (purple). The canonical interaction with the target (orange) occurs upstream of the boxes D and D’, using a region referred to as the ASE. The fifth nucleotide upstream of these boxes is methylated by the core C/D snoRNA interactor FBL. B) Schematic representation of experimentally validated noncanonical interaction regions. The classical elements are represented: the boxes C and C’ (green), D and D’ (purple) and the ASE (yellow). Bipartite interactions are represented by a dotted line. The names of the interacting snoRNA and target gene are indicated on the right. The reference for each interaction is indicated in superscript. The color of the interaction represents its effect, whether regulation of alternative splicing, regulation of RNA level (either pre-mRNA or mRNA) and RNA modification including methylation of noncanonical targets, acetylation and A to I editing. Combined together, noncanonical interactions cover the whole snoRNA.

In recent years, a wide range of functions have been discovered for box C/D snoRNAs, including the regulation of chromatin compaction, metabolic stress, cholesterol trafficking, alternative splicing and mRNA levels (reviewed in (9–11)). These functions are often mediated by noncanonical pairing configurations with the interactions involving diverse regions of the snoRNA sequence (Figure 1B) (8, 12–20). As such, complementarity to the ASE employed for methylation cannot be used as sole indicator of potential noncanonical targets. In this paper, we define a canonical interaction as an interaction leading to the methylation of a rRNA or a snRNA, and all others are defined as noncanonical, including interactions leading to the methylation of other types of RNA such as mRNA and transfer RNA (tRNA).

Genome wide methods for detecting RNA-RNA interactions identified a large number of noncanonical snoRNA interactions. These methods, including PARIS (21), LIGR-seq (14) and SPLASH (22), have been devised to survey all RNA duplexes in cells, both intra- and intermolecular. They have enabled the detection of known and novel snoRNA interactions. For example, LIGR-seq identified functional interactions between orphan snoRNA SNORD83B and three different mRNAs affecting their stability (14). Indeed, these large-scale experimental approaches play an important role in uncovering noncanonical snoRNA targets. However, these methods are limited by cell type, growth conditions and cross-linking approach used to produce the data. In addition, they often suffer from low proportion of intermolecular duplex reads (23) and consequently reduced coverage of snoRNA interactions, which form a relatively small proportion of all possible interactions in the human transcriptome, suggesting that there are probably plenty of snoRNA-RNA interactions yet to uncover.

In silico prediction of snoRNA interactions has the potential to uncover all possible transcriptome-wide snoRNA interactions since they are not restricted by experimental constraints. However, available RNA-RNA interaction prediction tools are often built on known mechanisms of function and interaction rules. For example, the C/D snoRNA interaction predictors, PLEXY (24) and snoscan (25), were developed to predict potential targets that interact with the canonical ASE sequence commonly used for rRNA modification. Indeed, PLEXY improves the efficiency of detecting canonical binding modes by only considering the 20 nucleotides upstream of the boxes D and D’ of the snoRNA, corresponding to the ASE, and filters the interactions using previously identified pairing constraints (7). Snoscan takes into account snoRNA features such as the boxes, the ASE, and the position between each element, but only considers the pairing in the 25 nucleotides upstream the boxes D and D’. Accordingly, while PLEXY and snoscan are efficient in identifying methylation targets, they have a limited use for identifying new forms of snoRNA-RNA interactions, especially those using an ASE independent homing sequence.

To help uncover a broader regulatory spectrum of box C/D snoRNAs and identify noncanonical interactions we developed snoGloBe, a C/D snoRNA interaction predictor based on a gradient boosting classifier. SnoGloBe considers all possible interactions between the snoRNA and targets regardless of the position within the snoRNA sequence. The predictor was trained and tested using known canonical interactions, experimentally detected large-scale snoRNA-RNA interaction datasets and validated noncanonical interactions from the human transcriptome. The accuracy and breadth of snoGloBe are evident from its capacity to recover most known snoRNA-RNA interactions, both canonical and noncanonical. Applying snoGloBe to the human coding transcriptome revealed positional enrichment of C/D snoRNA interactions in targets and functional enrichment for specific snoRNAs. The depletion of a model snoRNA altered the RNA level and splicing of a subset of the predicted targets. Notably, snoGloBe was also able to identify targets in a viral genome including a known interaction in the SARS-CoV-2 transcriptome. Overall snoGloBe provides a flexible discovery tool for box C/D snoRNA interactions that transcends pre-established binding rules.

## MATERIAL AND METHODS

### High-throughput RNA-RNA interaction analysis

The high-throughput RNA-RNA interaction (HTRRI) datasets from PARIS (21) (SRR2814761, SRR2814762, SRR2814763, SRR2814764 and SRR2814765), LIGR-seq (14) (SRR3361013 and SRR3361017) and SPLASH (22) (SRR3404924, SRR3404925, SRR3404936 and SRR3404937) were obtained from the short read archive SRA (26) using fastq-dump from the SRA toolkit (v2.8.2). The PARIS datasets were trimmed using the icSHAPE pipeline available at https://github.com/qczhang/icSHAPE. PCR duplicates were removed from LIGR-seq datasets using the script readCollapse.pl from the icSHAPE pipeline and the reads were trimmed using Trimmomatic version 0.35 with the following options : HEADCROP:5 ILLUMINACLIP:TruSeq3-SE.fa:2:30:4 TRAILING:20 MINLEN:25 (27). The quality of the reads was assessed using FastQC (v0.11.15) before and after the pre-processing steps (28). All the samples were analyzed using the PARIS pipeline as described in sections 3.7 and 3.8 from (29). Some modifications were made to the duplex identification and annotation scripts. The modified scripts are available at <u>github.com/Gabrielle-DF/paris</u>.

The RNA duplexes were assigned to genes using the annotation file described in (30) to which missing rRNA annotations from RefSeq (31) were added. The annotation file was modified using CoCo correct_annotation (32) to ensure the correct identification of snoRNA interactions.

Only the interactions between a box C/D snoRNA and a known gene were kept. To avoid intramolecular interactions, we removed interactions between a snoRNA and its 50 flanking nucleotides and interactions between two snoRNAs from the same Rfam family (33). To limit the number of false positives, we filtered the interactions based on their pairing using RNAplex (34). Only paired regions of the interactions were kept to get rid of unpaired flanking regions. The interactions were split at each bulge to ensure the correct alignment of the snoRNA and target sequences. The interactions shorter than 13 nucleotides were removed to match the length of the windows (see Input features section). We finally removed interactions that were already known and present in our positive set described in the next section. We obtained 445 box C/D snoRNA interactions (Figure S1).

### Positive set composition

The positive set is composed of the previously detected snoRNA interactions from PARIS, LIGR-seq and SPLASH filtered as described above, as well as interactions obtained from snoRNABase (35) and manually curated interactions from the literature (Table S1) (Figure 2A, B). Interactions from snoRNABase and from the literature that were shorter than 13 nucleotides were padded by adding their flanking sequence to respect the length threshold.

**Figure 2.**
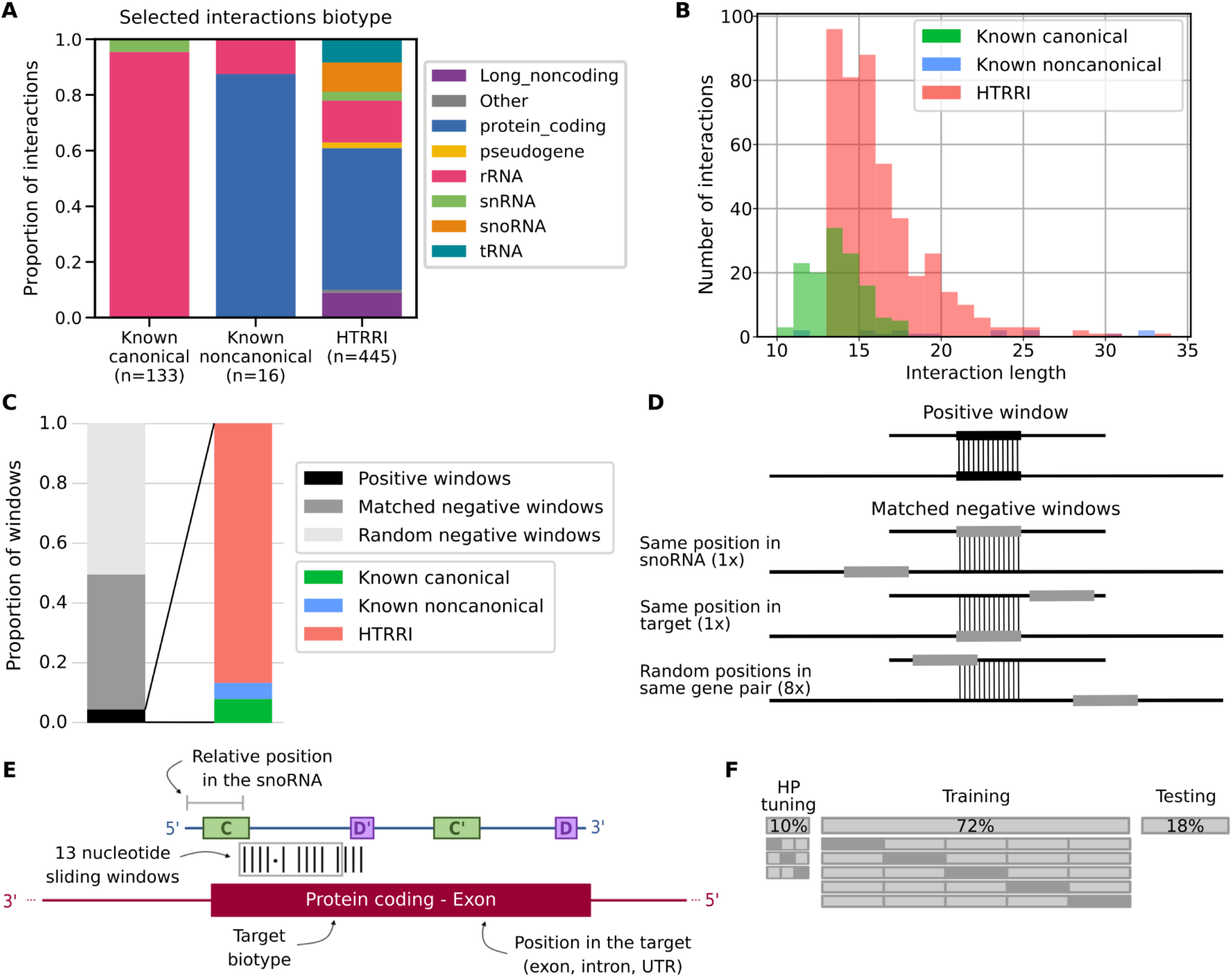
Composition of the dataset used to build snoGloBe. A) Diverse RNAs have been shown to bind box C/D snoRNAs. Interactions involving box C/D snoRNAs were collected and assembled including known canonical interactions with rRNA and snRNA from snoRNABase, known noncanonical interactions curated from the literature and interactions extracted from HTRRI datasets. The proportion of interactions involving different RNAs of each biotype is shown for each interaction source. The color legend for RNA biotypes is shown on the right. (B) Distribution of the length of interactions from each data source. (C) Distribution of the datasets used to build snoGloBe. The dataset consists of positive, matched negatives and random negatives in a proportion of 21 negatives (10 matched and 11 random) for 1 positive window. The positive windows are composed of HTRRI (86.3 %), known canonical (8.5 %) and noncanonical (5.2 %) interactions. (D) Generation of matched negative windows. 10 matched negative windows are generated for each positive one. The matched negative windows originate from the same snoRNA– target gene pair as the positive window. One has the same position in the snoRNA and a different position in the same target, one has a different position in the snoRNA and the same position in the target, and 8 windows have random positions in the same snoRNA-target pair. E) SnoRNA-RNA pairs are encoded for presentation to the predictor. Features considered include the 13 nucleotide sequence of the snoRNA and the 13 nucleotide sequence of the target, the relative position of the window in the snoRNA, the target biotype and the position in the target. F) The dataset is split in non-overlapping sets for hyperparameter tuning (10% of the windows), training (72% of the windows) and testing (18% of the windows). The hyperparameter tuning was done using a random search with 3-fold cross-validation. The model was trained and evaluated using stratified 5-fold cross-validation to ensure the correct representation of each category of positive windows in each subset. The known noncanonical windows were all kept for the validation set.

### Negative set composition

The negative set is composed of random negatives and matched negatives (Figure 2C). The random negative examples are the combination of random sequences from any box C/D snoRNA and any gene, whereas the matched negative examples are random sequences coming from a positive snoRNA-target gene combination (Figure 2D). We created 21 negative examples per positive example to reflect the fact that the majority of the transcriptome is not bound by snoRNAs. This negative set was split in three for hyperparameter tuning, training and testing as described in the following Redundancy removal for tuning, training and test sets section. As an additional validation step, we added seven negative windows for each positive window of the test set obtained from dinucleotide shuffle using the algorithm from Altschul and Erikson (36) implemented by P. Clote (http://clavius.bc.edu/~clotelab/RNAdinucleotideShuffle/ShuffleCodeParts/altschulEriksonDinuclShuffle.txt). We also added random negative windows to obtain a 1000 random negative examples for each positive example, amplifying the class imbalance to simulate the transcriptome.

### Input features

The interactions obtained from snoRNABase, the literature and HTRRI methodologies as described in the positive set composition section were split in 13 nucleotide sliding windows, with a step of 1 nucleotide. The 13 nucleotide window length was chosen to limit the chance of finding this sequence randomly in the genome, and most of the known interactions meet this length criteria (Figure 2B). Indeed, 13 nucleotides is rather small considering the size of the transcriptome, but the majority of known canonical interactions are longer or equal to this length (Figure 2B) and using a longer window would result in the loss of many validated windows or in the addition of flanking nucleotides, adding noise to the validated interaction set. Also, the sequence is only one part of the features used by the model to classify the interactions, so the small window size is compensated by the complementary information, such as the position in the snoRNA. The position of the interaction in the snoRNA varies depending on the interaction type. The canonical interactions are located in the ASE, whereas other interactions are distributed throughout the snoRNA (Figure 1B). We thus included the relative position in the snoRNA as an input feature. The information relative to the target biotype and the interaction position in the target were also used as input to account for the vast diversity of snoRNA interactors (Figure 2A) and gain insight into potential snoRNA function. In summary, each interaction window is composed of 13 nucleotides of the snoRNA and the corresponding 13 nucleotides of the target. The input features used are the window sequences in one-hot encoding, the relative position in the snoRNA between 0 and 1, the location in the target gene (intron, exon, 3’UTR and/or 5’UTR) and the target biotype (Figure 2E). The biotypes considered are listed in snoGloBe’s manual. Protein coding, pseudogene and long noncoding RNA biotypes were grouped according to http://vega.archive.ensembl.org/info/about/gene_and_transcript_types.html and http://ensembl.org/Help/Faq?id=468.

### Redundancy removal for tuning, training and test sets

The positive and negative examples were split into hyperparameter tuning, training and test sets. First, to remove redundancy from the sets, the snoRNAs were grouped based on their Rfam identifier (33). To ensure that the model is not trained and tested on similar snoRNAs, the Rfam families and clans were distributed in order to assign 18% of all examples in the test sets, with similar proportion of the initial HTRRI and known canonical interactions. We ensured that all members of a clan (or a family if the family is not part of a clan) are entirely included either in the training or the test set, but not both. All the known noncanonical interactions were kept for the test sets since there are very few such examples. The remaining examples were split between the hyperparameter tuning and training sets to consist respectively of 10% and 72% of initial data (Figure 2F). The interactions were split only based on the snoRNA space, and not the target. Since most snoRNAs target rRNA, it would be near impossible to form sets with interactions from distinct snoRNA families and clans, distinct targets, and having a representative mix of canonical and noncanonical interactions in all sets.

### Building the model

The model used is a gradient boosting classifier from scikit-learn (v0.21.3) (37). We selected the hyperparameters using a random search with 3-fold cross-validation due to the small number of interaction windows in the tuning set. The hyperparameters that were tuned are the number of trees (n_estimator), the minimum number of samples in a node to be split into two branches (min_samples_split), the minimum number of samples in a leaf (min_samples_leaf), the maximum depth of each tree (max_depth) and the value by which the contribution of each tree is decreased (learning_rate). The hyperparameter values selected by random search are: n_estimators = 371, min_samples_split = 76, min_samples_leaf = 49, max_depth = 2 and learning_rate = 0.43, others are kept to default.

The model was trained on the whole training set and validated with 5-fold stratified cross-validation, to keep similar proportions of HTRRI and known canonical interactions in each subset. A 5-fold cross-validation was used in the training validation instead of 3-fold cross-validation as the hyperparameter tuning step. The greater number of examples in the training set allowed to create more subsets having a good representation of each interaction category, and more validation steps increasing the confidence in the resulting model, called snoGloBe.

The model performance was evaluated on the test set and compared to PLEXY (24), snoscan (25), RNAplex (34), RIsearch2 (38), IntaRNA (39) and RNAup (34). To compare their performance on a similar basis, we selected a threshold (either a score for snoGloBe and snoscan, or an energetic cut-off for the other tools) resulting in 95% precision on the test set.. The details are available in Figure S2. PLEXY was only used for snoRNAs with non-degenerated boxes D and D’ to avoid bias caused by misidentified boxes.

### Prediction against protein coding genes

SnoGloBe was used to predict box C/D snoRNA interactions with protein coding transcripts. For this analysis, only box C/D snoRNAs expressed at 1 transcript per million (TPM) or more in at least one of the RNA-seq datasets from 7 different healthy human tissues (3 samples from different individuals for each of the following tissues: brain, breast, liver, ovary, prostate, skeletal muscle and testis) from a previous study (40) (available from GEO (41): GSE126797, GSE157846) were considered, totaling 312 snoRNAs. We predicted the interactions of these snoRNAs against all protein coding genes, split in 13-nucleotide windows with a step of two. We took whole gene sequences to predict interactions with any intron and exon. To narrow the number of predictions obtained, we kept the interactions having at least three consecutive windows with a score >= 0.98 for further analysis. The gene ontology enrichment analysis of the predicted targets was done using g:Profiler with all protein coding genes used for the interaction prediction as background (42).

### Overlap between predicted snoRNA interactions and eCLIP regions

All the eCLIP datasets (43, 44) were downloaded from the ENCODE portal (45), totaling 225 samples considering 150 proteins. The complete list of the datasets is available in Table S2. Only the eCLIP regions having a p-value <= 0.01 were kept. Datasets from the same protein were merged using BEDTools merge -s (v2.26.0) (46). The number of overlaps between the predicted interactions and the eCLIP regions was computed using BEDTools intersect -s.

### SNORD126 knockdown

HepG2 cells were cultured in complete Eagle’s Minimum Essential Medium (EMEM from Wisent) and passaged twice a week, according to ATCC guidelines. Trypsinized cells were then seeded at 350000 cells/well in 6 well plates in 1ml EMEM. Cells were transfected 24 hours later with 2 different antisense oligonucleotides (ASOs) targeting SNORD126 (30nM or 40nM) using Lipofectamine 2000 (LIFE technologies) and optiMEM (Wisent). A scrambled ASO was used as a negative control. The sequence of the ASOs are listed in Supplemental Figure S3.

Cells were harvested 48 hours post transfection, washed and pelleted, then resuspended in 1ml Trizol and stored at -80℃ until RNA extraction. This was repeated 3 times to obtain biological triplicates.

### RNA extraction

Total RNA extraction from transfected HepG2 cells was performed using RNeasy mini kit (Qiagen) as recommended by the manufacturer including on-column DNase digestion with RNase-Free DNase Set (Qiagen). However, 1.5 volumes Ethanol 100% was used instead of the recommended 1 volume ethanol 70% in order to retain smaller RNA. RNA integrity of each sample was assessed with an Agilent 2100 Bioanalyzer. RNA was reversed transcribed using Transcriptor reverse transcriptase (Roche) and knockdown levels were evaluated by qPCR.

### RNA-seq library preparation and sequencing

RNAseq libraries were generated from 1ug DNA-free total RNA/condition using the NEBNext® Ultra™ II Directional RNA Library Prep Kit for Illumina (E7760S) and following the Protocol for use with NEBNext Poly(A) mRNA Magnetic Isolation Module (NEB #E7490). The resulting libraries were submitted to a total of 10 cycles of amplification then purified using 0.9X Ampure XP beads. Quality and size was assessed with an Agilent 2100 Bioanalyser. Libraries were then quantified using a Qubit fluorometer, pooled at equimolar concentration and 1.8pM was sequenced on Illumina’s NextSeq 500 using a NextSeq 500/550 High Output Kit v2.5 (150 cycles) paired-end 2x75bp.

### RNA-seq analysis

The resulting base calls were converted to fastq files using bcl2fastq v2.20 (Illumina) with the following options: --minimum-trimmed-read-length 13, --mask-short-adapter-reads 13, --no-lane-splitting. The fastq files were trimmed for quality and to remove remaining adapters using Trimmomatic v0.36 (27) with ILLUMINACLIP:{adapter_fasta}:2:12:10:8:true, TRAILING:30, LEADING:30, MINLEN:3. The sequence quality was assessed using FastQC v0.11.5 (28) before and after the trimming. The trimmed sequences were aligned to the human genome (hg38) using STAR v2.6.1a (47) with the options --outFilterScoreMinOverLread 0.3, -- outFilterMatchNminOverLread 0.3, --outFilterMultimapNmax 100, -- winAnchorMultimapNmax 10, --alignEndsProtrude 5 ConcordantPair. Only primary alignments were kept using samtools view -F 256 (v1.5) (48). The gene quantification was done using CoCo correct_count -s 2 -p (v0.2.5p1) (32). DESeq2 was used for the differential expression analysis (49). Genes having a corrected p-value <= 0.01 were considered significantly differentially expressed. The alternative splicing analysis was done using MAJIQ v2.2 and VOILA with the option --threshold 0.1 (50).

### Empirical p-value calculation

To evaluate the significance of the overlaps between each snoRNA predicted interaction and eCLIP binding sites, alternative splicing events and differentially expressed genes, we computed an empirical p-value using BEDTools shuffle 10 to 100 000 times for each combination followed by BEDTools intersect -s through pybedtools (46, 51). BEDTools shuffle was used with an appropriate background for each analysis: all protein coding genes for eCLIP binding sites and protein coding genes having an average of 1 TPM across all sequencing datasets for differential expression and alternative splicing analyses. We counted the number of distinct events, or genes in the case of differential expression analysis, having an overlap with at least one shuffled interaction for each iteration. P-values were calculated as the proportion of iterations in which the shuffled dataset overlap was at least as extreme as the true dataset overlap.

### Prediction of human snoRNA interaction with SARS-CoV-2 transcriptome

We predicted the interactions between the expressed human snoRNAs against SARS-CoV-2 transcriptome, using SARS-CoV-2 ASM985889v3 genome assembly and the annotation file Sars_cov_2.ASM985889v3.101.gtf obtained from Ensembl COVID-19 (52). We used thresholds of a minimum of 3 consecutive windows having a probability greater or equal to 0.85.

### Reagents

The following reagents were used in this study. Eagle’s Minimum Essential Medium (purchased from Wisent located at St-Bruno, Québec, Canada, catalog number 320-005-CL) and optiMEM (purchased from Wisent located at St-Bruno, Québec, Canada, catalog number 31985-070) were used for tissue culture, lipofectamine 2000 (LIFE technologies, Burlington, Ontario, Canada, catalog number 11668019) for transfection, RNEasy mini kit (Qiagen, Mississauga, Ontario, Canada, Catalog number 74106) and RNase-Free DNase set (Qiagen, Mississauga, Ontario, Canada, catalog number 79254) both for RNA extraction, Transcriptor reverse transcriptase (Roche, Laval, Québec, Canada, catalog number 3531287001) for RNA reverse transcription for RT-PCR, NEBNext® Ultra™ II Directional RNA library Prep Kit (New England Biolabs, Pickering, Ontario, Canada, catalog number E7760S) and NEBNext Poly(A) mRNA Magnetic Isolation Module (New England Biolabs, Pickering, Ontario, Canada, catalog number E7490) for library preparation, NextSeq 500/550 High Output Kit v2.5 150 cycles (Illumina, Vancouver, British Columbia, Canada, catalog number 20024907) for sequencing.

### Biological Resources

The HepG2 cell line used for SNORD126 knockdown was obtained from ATCC (HB-8065). The antisense oligonucleotides used for the knockdown were purchased from IDT and their sequences are listed in Figure S3.

### Statistical Analyses

The empirical p-values were calculated as described above. The p-values of the overlap between eCLIP binding sites and snoRNA interaction predictions were adjusted according to Bonferroni correction by multiplying the empirical p-value obtained by 150, the number of RBP tested for each snoRNA. SNORD126 knockdown was done using 2 ASOs and 1 control ASO, each having four biological replicates. The adjusted p-values for the differential expression analysis were obtained by DESeq2, which applies the Benjamini-Hochberg correction. The equations used to evaluate the precision, recall, false positive rate, accuracy, F_1_ score and Matthews correlation coefficient shown in Figure 3 are listed below. For snoGloBe, RNAup, RNAplex, IntaRNA and RIsearch2, an interaction window was deemed positive if the score between the two 13-nucleotide windows was greater than the threshold specified in Figure S2A in absolute terms. For PLEXY and snoscan, the full snoRNA must be given as input, so, to ensure fairness when comparing to other tools, the interaction windows were considered positive if the predicted interaction occurred inside the 13-nucleotide region of the snoRNA that was considered for this target window and the score was greater than the threshold in absolute terms, the interactions predicted outside this region were ignored (Figure S2B). The interaction windows were then classified as true positive (TP), true negative (TN), false positive (FP) or false negative (FN) depending their predicted status described above and to which set (positive or negative) they belonged.

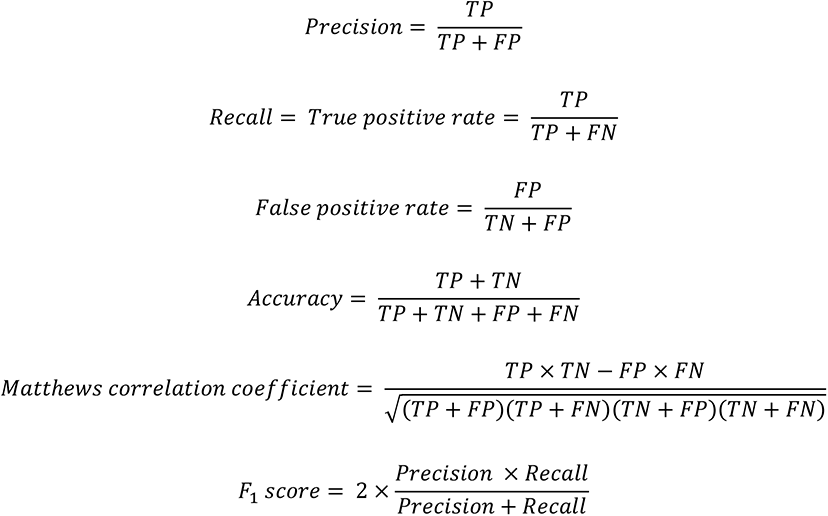

**Figure 3.**
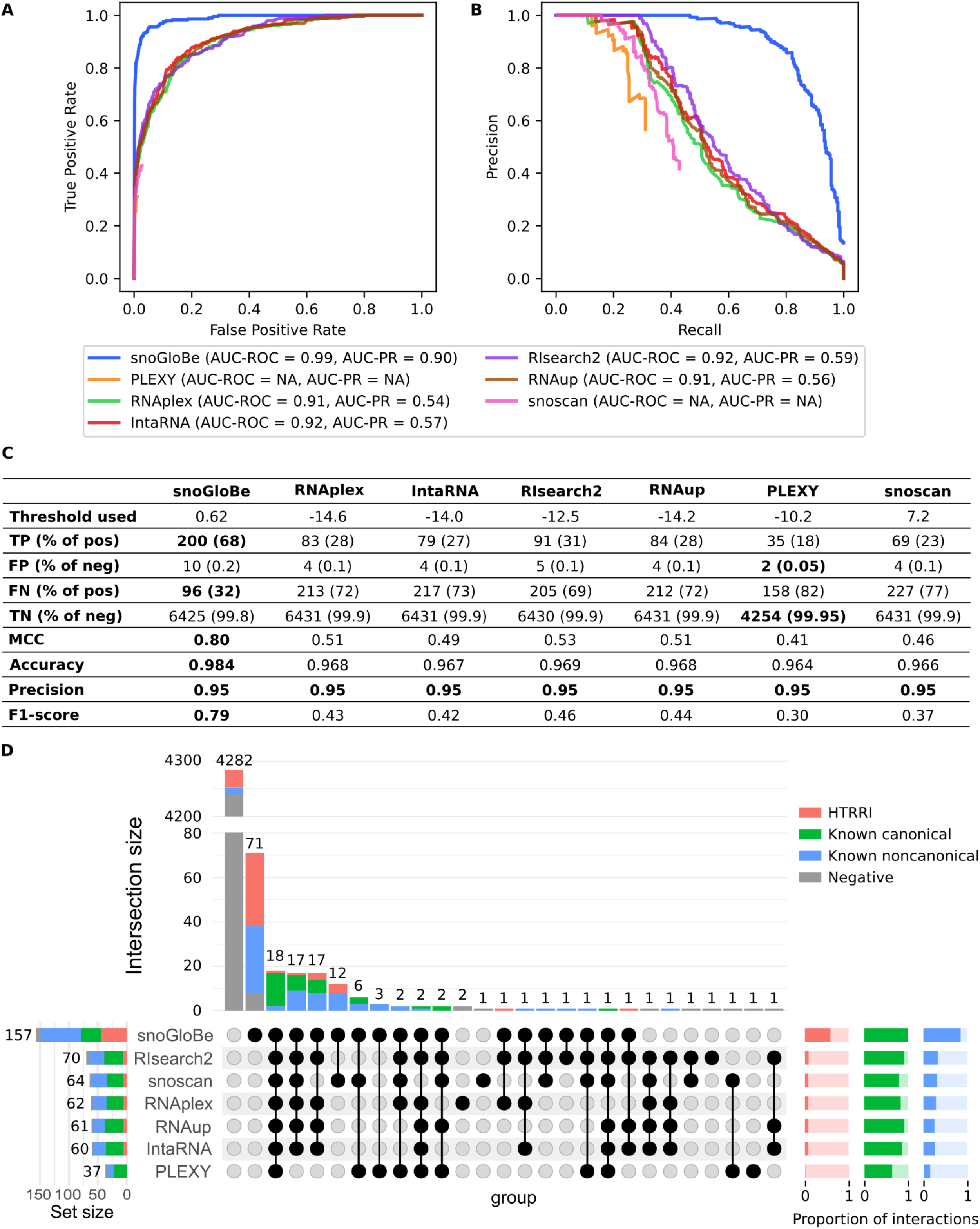
SnoGloBe performs better than available tools. (A) Receiver Operator Characteristic (ROC) and (B) Precision-Recall (PR) curves of different tools calculated on the test set. The corresponding area under the curves (AUC) are indicated in the legend. C) Table of performance measures from different tools calculated on the test set with a threshold set to obtain a precision of 95%. PLEXY was only used on interactions from box C/D snoRNA with non-degenerated boxes D and D’ (Table S1) since the position and sequence of the boxes are required. D) Upset plot representing the overlaps between each tool prediction of the test set’s positive windows. The upset plot only shows the subset of the interactions that were considered for PLEXY to ensure a fair comparison. The upset plot of all the test set’s positive windows is shown in Figure S8. The proportion of positive examples from each category predicted as positive (dark color) and negative (light color) by each tool is shown in the bottom right.

### Data Availability/Sequence Data Resources

All raw and processed sequencing data generated in this study have been submitted to GEO (41) under accession number GSE184173. The datasets used to evaluate snoRNA expression are available under the accession numbers GSE126797 and GSE157846. HTRRI datasets used for building snoGloBe are available from SRA (26) under the following accession numbers : SRR2814761, SRR2814762, SRR2814763, SRR2814764, SRR2814765, SRR3361013, SRR3361017, SRR3404924, SRR3404925, SRR3404936 and SRR3404937.

### Data Availability/Novel Programs, Software, Algorithms

The snoGloBe interaction predictor described in this paper is available from https://github.com/scottgroup/snoGloBe. The modified PARIS (29) scripts for the analysis of high-throughput RNA-RNA interaction datasets are available at <u>github.com/Gabrielle-</u> <u>DF/paris</u>.

### Web Sites/Data Base Referencing

The following tools and websites were used in this study: bcl2fastq v2.20 (https://support.illumina.com/sequencing/sequencing_software/bcl2fastq-conversion-software.html), FastQC v0.11.5 (28), Trimmomatic v0.36 (27), STAR v2.6.1a (47), CoCo v0.2.5p1 (32), SAMtools v1.5 (48), BEDTools v2.26.0 (46), pybedtools v0.8.1 (51), MAJIQ v2.2 (50), VOILA v2.2.0-e25c4ac (50), DESeq2 (49), scikit-learn v0.21.3 (37), PLEXY (24), RNAplex v2.4.14 (34), RIsearch2 v2.1 (38), IntaRNA v3.1.1 (39), RNAup v2.4.14 (34), RefSeq (https://www.ncbi.nlm.nih.gov/refseq/) (31), Ensembl (https://www.ensembl.org/) (52), ENCODE portal (https://www.encodeproject.org/) (45), GEO (https://www.ncbi.nlm.nih.gov/geo/) (41), SRA (https://www.ncbi.nlm.nih.gov/sra) (26), SRA Toolkit v2.8.2 (https://github.com/ncbi/sra-tools), snoRNABase (https://www-snorna.biotoul.fr/) (35), icSHAPE (https://github.com/qczhang/icSHAPE).

## RESULTS

### Identification and curation of experimentally identified snoRNA-RNA interactions

In order to create a predictor of snoRNA-RNA interactions, we began by extracting the canonical snoRNA interactions from snoRNABase (35) and we combined it with experimentally validated noncanonical interactions reported in the literature. As a result, we identified 149 non-redundant interactions, 133 from snoRNABase and 16 from the literature (Figure 2A). In addition, we performed a de novo analysis of the datasets obtained from three high-throughput RNA-RNA interaction (HTRRI) identification methodologies PARIS (21), LIGR-seq (14) and SPLASH (22) to extract new experimentally detected interactions. All sets were analyzed using the PARIS bioinformatics protocol (29), filtered to remove interactions shorter than 13 base pairs and the interactions featuring bulges were split to remove the unpaired region (Figures 2B, S1). Since HTRRI are identified in a high-throughput manner and have no functional validation, some can be false positives solely occurring by chance and be inconsequential. This adds noise to the dataset and must be kept in mind throughout this study. Overall, we retrieved 133 known canonical interactions from snoRNABase, 16 known noncanonical interactions from the literature and 445 putative HTRRI, totalling 594 interactions listed in Table S1. The identified target biotypes varied greatly based on the data source underscoring the effect of the experimental approach and RNA source (Figure 2A). While snoRNABase is the main repository of canonical human snoRNA interactions, our manual curation of the literature revealed articles describing noncanonical snoRNA interactions and thus mainly involves protein coding targets. In contrast, PARIS, LIGR-seq and SPLASH are methodologies detecting RNA-RNA interactions with less experimenter bias. The widest distribution of target biotypes was identified in the HTRRI dataset and the largest number of distinct targets was found in protein coding RNAs (Figure 2A). The newly generated combined interaction set includes a wide variety of biotypes and interaction modes and forms an excellent base for a positive set. To create a negative dataset, we generated a combination of random negative and matched negative interactions (Figure 2C). The random negative examples are random sequence pairs from any box C/D snoRNA and any gene, whereas the matched negative examples are random sequences originating from a snoRNA-target gene combination from the positive set (Figure 2D). The positive:negative ratio was chosen to be imbalanced (1:21) to reflect the fact that the proportion of transcriptomic sequences bound by C/D snoRNAs is expected to be much lower than the proportion not bound.

### Feature encoding and predictor training

Since snoRNA-RNA interactions involve the formation of an RNA duplex, the sequences of the two RNAs must be encoded amongst the features presented in input. The duplex length of validated canonical and noncanonical snoRNA interactions varies from 10 to 32 base pairs (Figure 2B, known canonical and known noncanonical), so we encoded the interaction sequences in windows of 13 nucleotides for both the snoRNA and its target. This compromise helps us take into account the validated snoRNA-RNA interactions while limiting the chance of finding these sequences randomly in the genome. The position of the interaction window in the snoRNA varies greatly between the canonical and noncanonical interactions. Indeed, canonical interactions employ only the regions immediately upstream of the boxes D or D’ while the regions involved in noncanonical interactions cover the entire snoRNA length (Figure 1B). Therefore, we did not specify a fixed position but instead included it as an input feature. To gain information about the possible functional impact of the interactions we also included the target biotype as well as the position in the target (either in an exon and/or an intron and whether the exon is a 5’ or 3’ UTR when appropriate) as input features (Figure 2E). These input features were encoded for all positive and negative snoRNA-RNA pairs. Since snoRNA-RNA duplexes are encoded as 13 nucleotide pairs of windows, an interaction can consist of multiple such pairs of windows. The resulting datasets consist of 1740 positive and 36865 negative windows (Figure 2C). The datasets were split in a non-overlapping manner for hyperparameter tuning, model training and model testing in a 10:72:18 relative proportion (Figure 2F). However, since there are few examples of known noncanonical interactions, they were all kept for the test set (Table S1). To avoid the redundancy caused by multicopy snoRNA genes (53, 54) we made sure that members of the same snoRNA clan (or for snoRNAs that are not part of a clan, we considered families, as defined for both clans and families by Rfam (33)) are present in only one dataset. For example, no member of the SNORD33 clan is included in the test set if other members of the family are present in the training set (details in Methods section). Together the selection criteria allow the identification of a broad range of targets, reduce redundancy and decrease the dependency on the canonical mode of interaction.

### SnoGloBe accurately predicts a wide range of interactions

The model used by snoGloBe is a gradient boosting classifier, which is a combination of multiple decision trees. The hyperparameters were tuned on 10% of the data, using a random search. The model was then trained on 72% of the data using a 5-fold stratified cross-validation (Figure 2F). The output of the prediction is a value between 0 and 1 representing the probability of interaction. The performance of the model was then evaluated on the test set. As indicated in Figure S4, snoGloBe clearly separated the negative and positive examples in the independent test set, giving the great majority of the negatives (96%) a score below 0.1 and the majority of positives (63%) a score above 0.9.

To our knowledge, there is currently no available snoRNA target prediction tool that is built to predict both canonical and noncanonical box C/D snoRNA interactions, so the model was compared to the closest tools we could find: snoRNA specific interaction predictors and general RNA duplex predictors including PLEXY (24), snoscan (25), RNAup (34), RNAplex (34), RIsearch2 (38) and IntaRNA (39) using the parameter values summarized in Figure S2. Since these tools were not created for this specific task, they are not expected to perform as well, especially in the case of canonical snoRNA interaction predictors that were trained to classify the noncanonical interactions as negatives. The goal of this comparison is to show that snoGloBe is a good addition to the current available tools, and not an evaluation of the other tools’ ability to perform the task for which they were designed. The comparison of these tools on the test set shows that snoGloBe outperformed the other tools in predicting the snoRNA interactions by obtaining the highest area under the ROC and precision-recall curves (Figure 3A, B). In addition, all the tools performed similarly on the test set and the training set, including snoGloBe, hinting that the model is able to generalize its learning on examples it has never seen before (compare Figure S5 and Figure 3A-B). PLEXY has the weakest performance, which is expected since it only predicts interactions with the ASE and the test set has interactions with all regions of the snoRNA (Figure S6). Interestingly, snoGloBe performs better than general RNA-RNA interaction predictors, hinting that snoGloBe is doing more than simple base-pair matching by capturing the specific information defining snoRNA - target interactions. Indeed, an analysis of the importance of each feature shows that the relative position of the interaction in the snoRNA has a major role in the classification, followed by information regarding the sequence of the interaction in the snoRNA and the target, and some biotypes, such as pseudogene and rRNA (Figure S7). This indicates that snoGloBe takes into account a variety of information to classify the interactions and not only the base-pairing.

We then determined the number of test set windows from each category predicted as positive and negative for each tool by using a threshold to obtain a 95% precision with every tool (Figure S2). SnoGloBe retrieves the highest number of true positive windows, and the highest proportion of known canonical, known noncanonical interactions and HTRRI (Figures 3C, D, S8). PLEXY retrieves the smallest number of positive windows from the test set, and most are from known canonical interactions. On the other hand, snoscan performs similarly to generic RNA-RNA interaction prediction tools and is able to retrieve interactions from all three categories, but its performance decreases with the addition of snoRNAs with degenerated boxes D and/or D’ (compare Figures 3D and S8). Generic RNA-RNA interaction predictors give similar results amongst themselves and retrieve mostly known canonical and noncanonical interactions. Interestingly, although snoGloBe was not trained on any known noncanonical interaction, it outperformed the other tools by identifying 74/94 noncanonical windows (Figure 3D, S8). Every tool tested was able to retrieve most of the known canonical interactions, but only a minority of known noncanonical interactions and even fewer HTRRI, except for snoGloBe which was able to predict the majority of all three categories of positive examples. The least well predicted interaction category is the HTRRI. Since the HTRRI identification methodology is prone to false positives, it is possible that the misclassified HTRRI are in fact interactions only happening by chance.

As an additional validation step, we added negative examples obtained by dinucleotide shuffling of positive interaction windows. These negative sequences differ from the positive examples, but all the other input features stay the same. As mentioned earlier, snoGloBe relies on a variety of input features to classify a potential interaction, so the shuffling of the sequence without modifying any other input features represents a greater challenge for snoGloBe than for the predictors mainly based on the sequence complementarity. We also added random negative windows to obtain a ratio of 1000 random negative for each positive example, to exacerbate the class imbalance and better represent the fact that transcriptome is mostly not bound by C/D snoRNAs. As shown in Figure S9A, every tool’s performance is affected by the addition of these negative examples, but snoGloBe still displays the highest area under the ROC and PR curves. The dinucleotide shuffled windows are the hardest negative examples to classify as they get the higher scores with every tool (Figure S9B). SnoGloBe still exhibits a good separation between the positive window scores and all three types of negative examples. Taken together, these data show that snoGloBe outperforms current predictors as it predicts more interactions with higher diversity and greater overall accuracy.

### Transcriptome-wide snoGloBe predictions reveal an enrichment of snoRNA interactions in messenger RNA regulatory sequences

The interactions of every expressed human snoRNA were predicted against all protein coding genes in human. As many snoRNA copies in human are not expressed and likely represent ‘dead’ copies in the genome (53), we restricted our study to only those detected as expressed as described in the Methods section. To limit the number of predicted interactions, we used a stringent cut-off of 3 consecutive windows having a probability (i.e. snoGloBe output score) greater than or equal to 0.98 (Figure S10). With this threshold, we obtain a median of 1017 predicted interactions per snoRNA (Figure 4A). SnoRNAs with the highest number of predicted interactions include snoRNAs with known validated noncanonical targets like SNORD32A (>1500 interactions), SNORD83B (>6000 interactions) and SNORD88C (>10 000 interactions) (8, 14, 15). The global analysis of all snoRNA predicted interactions reveals a preference for the region upstream of the box D, even though interactions were predicted throughout the whole snoRNA length and the training and test sets displayed enriched interactions upstream of both the boxes D and D’ (Figure 4B, Figure S6). Interestingly however, individual snoRNAs show different accumulation profiles along their length including some with a clear preference for targets binding the region upstream of the boxes D’ or D and others with a square accumulation in regions other than those upstream of boxes D/D’. For example, SNORD45C displays two strong regions of target binding, found respectively upstream of the boxes D’ and D (Figure S11A). On the other hand, SNORD11 has only one such region, upstream of the box D (Figure S11B). In contrast, SNORD31B in Figure S11 panel C and SNORD18A in panel D both have only one clear target binding region, overlapping and downstream of the box D’. Interestingly, while most snoRNA target sequences were located in introns, we detected enrichment in the exons and exon-intron junctions when compared to the distribution of these features in the transcriptome (Fig 4 compare panels C and D). Indeed, the protein coding transcriptome consists of 7.26% exonic sequences and 0.03% intron-exon junctions (measured in terms of 13 nt windows as described above), while 15.65% and 1.19% of the predicted snoRNA interactions are found in these regions respectively, representing increases of >2 fold in exonic sequences and 40 fold in intron-exon junctions. Many exonic snoRNA interactions are found in the 5’ or 3’ UTRs (Figure 4E, F), suggesting a role in regulating translation, transcript stability and/or 3’ end processing. Binding of snoRNAs to exonic sequence accumulates in precise regions or hot spots arguing for an organized binding program (Figure 4G). In addition, we found 15 snoRNAs that are predicted to interact with highly similar copies of themselves located at different genomic loci (i.e. the snoRNA and its copy are not located in the same host gene). These copies are encoded on the opposite strand of a protein coding gene, but none of these copies have a host gene on the same strand, therefore, they are unlikely to be expressed since most snoRNA depend on the transcription of their overlapping host gene to be produced (Figure S12). This hints at the possibility that a snoRNA could be retrotransposed antisense to a protein coding gene to act as a regulatory region, as already described for miRNA (55, 56). We conclude that binding of snoRNAs is not randomly distributed in the human transcriptome but targets specific regulatory elements and in particular those regulating splicing and translation.

**Figure 4.**
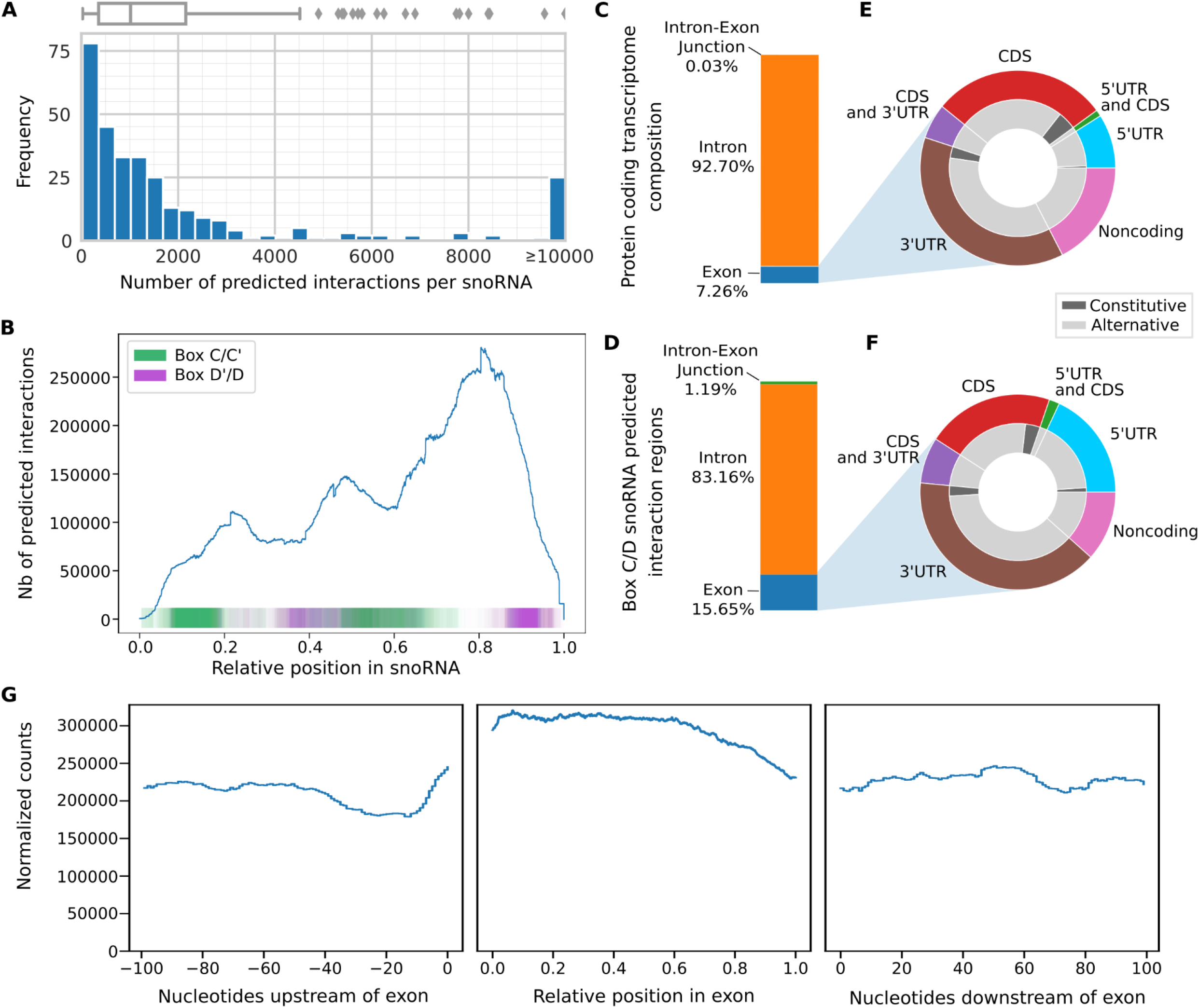
Box C/D snoRNA predicted interactions across the coding transcriptome. A) Histogram and boxplot (above) of the number of interactions per snoRNA using a threshold of at least 3 consecutive windows having a probability greater than or equal to 0.98. Most snoRNAs have less than 2000 predicted interactions. B) Distribution of the predicted region of interaction in all snoRNAs. The position in the snoRNAs is normalized between 0 and 1. The computationally identified boxes C, D’, C’ and D are respectively represented in green and purple. The predicted interactions are found throughout the snoRNA, with an enrichment in the 3’ end. C-D) Bar chart representing the proportion of exon, intron and intron-exon junction in the protein coding transcriptome (C) and the box C/D predicted interactions in the targets (D). The predicted interactions are enriched in the exons and the intron-exon junctions (D) compared to the protein coding transcriptome (C). E-F) Doughnut charts representing the composition in terms of 13-nt windows of the exons of the protein coding transcriptome (E) and the box C/D snoRNA predicted interactions in the targets (F). The predicted interactions located in exons are mainly found in UTRs (F) and are enriched in 5’UTRs when compared to the protein coding transcriptome (E). G) Distribution of the predicted interactions 100 nucleotides upstream of exons (left), in the exon (middle), and 100 nucleotides downstream of the exons (right). The positions in the exons are normalized between 0 and 1. The number of interactions is normalized by the number of existing features (exons or introns) at each position multiplied by one million. The predicted interactions are uniformly distributed across the exons, there is a higher number of interactions predicted inside the exons than in the flanking nucleotides.

### SnoGloBe uncovers functional specialization of snoRNA

Surprisingly, we found that snoRNAs do not feature uniform binding patterns or target preferences but instead display snoRNA or snoRNA family specific binding patterns. For example, SNORD35A binding is increased on the 3’ splice site, snoU2-30 binding is enriched in the 5’ splice site while SNORD38A shows an enrichment on the polypyrimidine tract (PPT) (Figure S13). Some snoRNAs display even more convincing target sets, including strong enrichment in specific gene elements of target genes enriched in specific biological processes, as well as significant overlap with functionally relevant RNA binding protein (RBP) target sites. For example, SNORD50B was found to be strongly enriched in 5’UTR binding of its targets and using its box D adjacent guide region (Figure S14A, B). Gene ontology enrichment analyses show that predicted targets of SNORD50B are involved in neuronal functionality and genes coding for proteins related to cell-cell interactions (Figure S14C). Many SNORD50B exonic targets bind alternative 5’UTRs, involving alternative transcription start sites, such as those of NDFIP2, COPS3 and SPG21, which could lead to transcript-specific effects (Figure S15). These possible regulatory events of SNORD50B are not randomly distributed but appear to target a specific group of genes. In contrast to SNORD50B which is specialized in targeting alternative 5’ UTRs, SNORD22 shows enriched binding at the 3’ splice sites and PPTs of its targets, involving a non-ASE region of the snoRNA overlapping with the box C’ (Figure S16A, B). Many SNORD22 3’ splice site targets bind alternatively spliced exons including in the diacylglycerol kinase zeta gene DGKZ, the amyloid beta precursor protein binding family B member APBB1 and three hits on the same alternatively spliced exon in the focal adhesion protein PXN (Figure S17). Gene ontology terms for SNORD22 targets are enriched in membrane proteins, cell junctions and GTPases (Figure S16C). Together these data indicate that snoRNAs use different mechanisms to identify their targets, co-regulating players of a common cellular function.

### Overlap of the binding sites of snoRNAs and RNA binding proteins

Comparison between the snoRNA binding sites identified by snoGloBe and the binding sites of RNA binding proteins (RBP) as determined by the ENCODE project using eCLIPs (43, 45) indicated strong overlap between the two. For example, both SNORD50B and SNORD22, which show strong position enrichment and functional enrichment of their targets, show strong overlap of their binding sites with those of specific RBPs (Figure S14D, S16D). In the case of SNORD50B, the strongest enrichments include DDX3X a helicase known to bind RNA G-quadruplexes in 5’UTRs, NCBP2 a cap-binding protein interacting with pre-mRNA, BUD13 involved in pre-mRNA splicing and FTO an RNA-demethylase involved in the maturation of mRNAs, tRNAs and snRNAs, supporting a role for SNORD50B in the maturation of pre-mRNA and in particular their 5’ extremity. In contrast, for SNORD22, the RBPs with strongest enrichment are PCBP2 the poly(rC) binding protein, PTBP1 a PPT binding protein involved in the regulation of alternative splicing as well as BUD13, PRPF8 and AQR all known as involved in pre-mRNA splicing, supporting a role for SNORD22 in the regulation of alternative splicing, through the binding of PPTs. These data suggest that snoRNAs may influence RBP function through collaborative or competitive binding to the targeted RNA sequence.

### SnoGloBe predicts functional regulatory targets

To evaluate the functional significance of snoGloBe’s predicted interactions, we experimentally measured the impact of knocking down a model snoRNA on its predicted targets. We chose SNORD126 as a model since it was implicated in different cellular functions while most of the targets relevant to these functions remain unidentified. This snoRNA was originally thought to be an orphan (35), but was later predicted to interact with and then shown to conditionally methylate the 28S rRNA (57, 58). In addition, it was shown that SNORD126 can activate the PI3K-AKT pathway through a yet to be determined mechanism (59). The SNORD126 interactions predicted by snoGloBe against protein coding genes led to interesting profiles. Most of the snoGloBe predicted interactions for this snoRNA involve a non ASE sequence overlapping the D’ box suggesting noncanonical methylation independent functions (Figure 5A and B). SNORD126 interactions are enriched in exons (1.6 fold) and particularly enriched on intron-exon junctions (>50 fold) compared to the protein coding transcriptome composition (Figure 5C-D). The exons predicted to be targeted by SNORD126 are enriched in 5’UTR (Figure 5E-F). SNORD126 predicted interactions are mostly uniformly distributed in the target exons, with the exception of notable enrichment around 80 nucleotides upstream of the exons (Figure 5G). These data suggest that SNORD126 may regulate gene expression through modulation of RNA stability and/or splicing.

**Figure 5.**
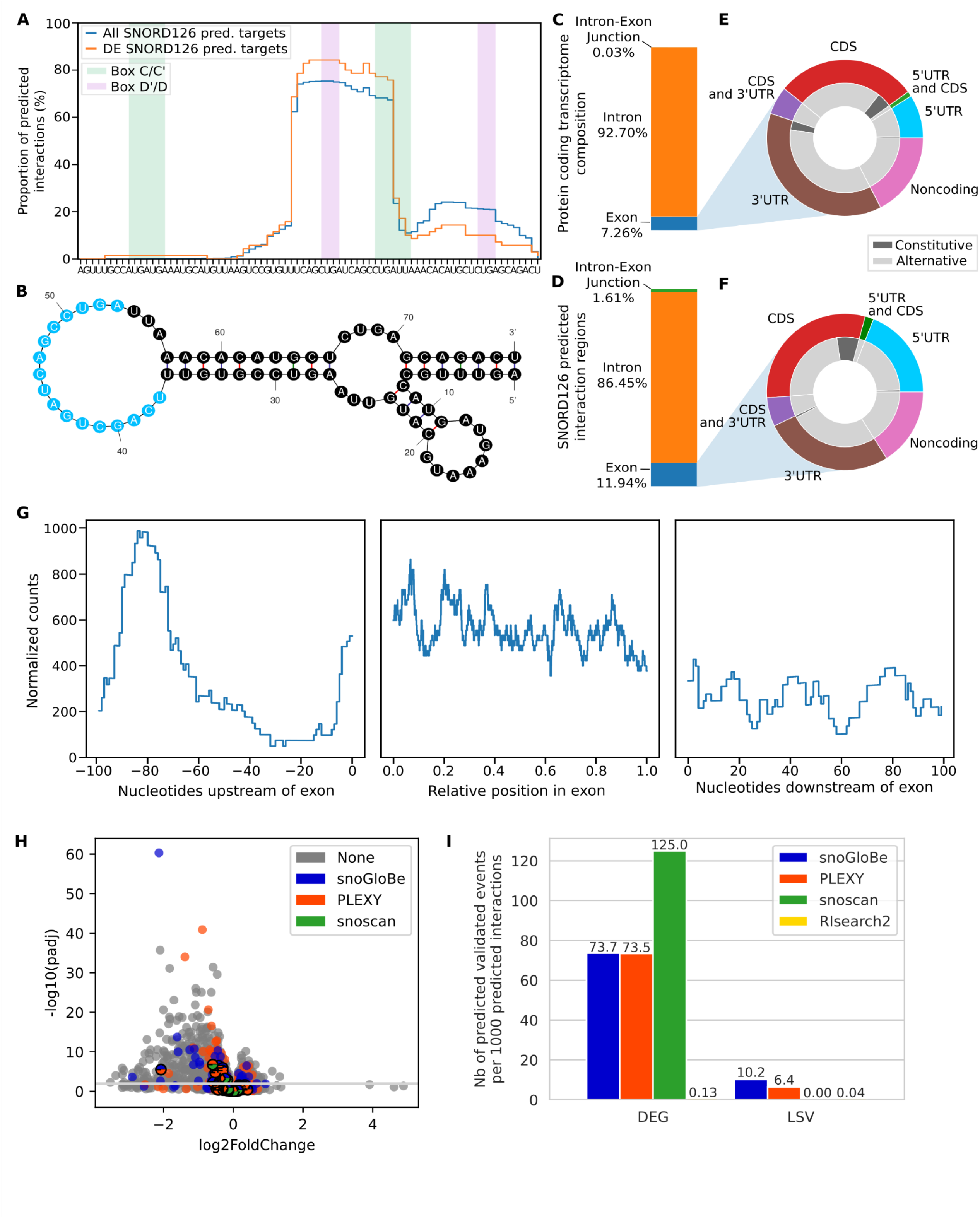
SNORD126 predicted targets are significantly affected by its knockdown. A) The major interaction site in SNORD126 predicted by snoGloBe is located in the middle of the snoRNA and doesn’t match the ASE upstream of the boxes D and D’ represented in purple. The accumulation profile represents the proportion of SNORD126 predicted interactions overlapping each nucleotide in the snoRNA (blue) and the interactions predicted in differentially expressed (DE) genes (orange). B) Predicted folded structure of SNORD126 considering the main region of interaction. Mfold (61) was used to predict the secondary structure of SNORD126, forcing nucleotides 37 to 53 to be single stranded (blue) through the unafold webserver. C-D) SNORD126 predicted interactions are enriched in the exons and the intron-exon junctions. (C) Shows the relative length of the different elements of the protein coding transcriptome while (D) shows the relative proportion of the targets of SNORD126. E-F) The predicted interactions located in exons are enriched in 5’UTRs. (E) Doughnut chart showing the breakdown of the different constituents of exons in the protein coding transcriptome. (F) Doughnut chart showing the same breakdown but only for regions targeted by SNORD126. G) SNORD126 predicted interactions are uniformly distributed across exons, with an enrichment around 80 nucleotides upstream of exons. H) Volcano plot representing the impact of SNORD126 knockdown on protein coding genes. The dots above the gray line are considered significantly differentially expressed (adjusted p-value ≤ 0.01). SNORD126 targets predicted by snoGloBe are colored in blue, PLEXY in orange and snoscan in green. Targets predicted by more than one tool are represented by dots colored by all corresponding tools. Only genes having a mean of 1 TPM across all samples are shown. (I) Bar plot representing the number of experimentally validated events (LSV : local splicing variation, DEG: differentially expressed gene) having at least one overlapping predicted interaction per thousand total SNORD126 predicted interactions by snoGloBe, PLEXY, snoscan and RIsearch2.

To evaluate the impact of SNORD126 on splicing and RNA stability we knocked it down using RNase H dependent antisense oligonucleotides and monitored the impact on the transcriptome using RNA-seq. The knockdown was performed in the HepG2 cell line, which is frequently used in genome wide analysis (45). In this cell line, 798 predicted target genes are expressed at 1 TPM or higher. Overall, the knockdown of SNORD126 resulted in the up regulation of 340 genes and the down regulation of 710 genes, totalling 1050 protein coding genes, 65 of which are snoGloBe’s predicted targets (Figure 5H). The overlap between the predicted targets and the differentially expressed genes shows a significant enrichment (p-value < 0.0001 by random sampling analysis as described in the Methods). The most upregulated predicted target following SNORD126 knockdown is BNIP3L, a pro-apoptotic protein, and the predicted interaction is located in the intron. The upregulation of BNIP3L following SNORD126 knockdown is in line with SNORD126 oncogenic role through the activation of PI3K-AKT pathway (59) and this predicted interaction could be an interesting lead to elucidate the underlying mechanism. On the other hand, the most downregulated predicted target is DNAH17, a dynein component, and has two predicted binding sites, one in an exon and one in an intron, around 300 nucleotides upstream the exon. These data suggest that one snoRNA can have different effect depending on the binding characteristics, such as the number of interactions and their region.

In addition to the change in expression, SNORD126 knockdown also altered the alternative splicing of transcripts. 309 such events are affected by the knockdown of SNORD126, 9 of which overlap a predicted SNORD126 binding site (p-value = 0.002). Amongst the interesting candidates, the target site of SNORD126 on CPT1B, encoding a carnitine O-palmoyltransferase, overlaps an alternative 5’ splice site detected with a differential splicing pattern and the target site of SNORD126 on MR1, which encodes a hydrolase involved in the NF-kB pathway, is near a 3’ splice site for which the intron has an alternative 5’ extremity (Figure S18). Interestingly, three genes having an alternative splicing event overlapping a predicted interaction were also differentially expressed upon SNORD126 knockdown: CPT1B, MR1 and DDX11, hinting that SNORD126 could affect RNA stability through alternative splicing.

To further evaluate snoGloBe’s performance in predicting functionally relevant interactions, we compared it to PLEXY (24), snoscan (25) and RIsearch2 (38), the general RNA-RNA interaction predictor with the best area under the ROC and Precision-Recall curves on the test set (Figure 3A, B). For PLEXY and RISearch2, we used an energy threshold of -20.4 kcal/mol, which is the average of snoRNA-rRNA duplexes recovered by PLEXY (24), and for snoscan we used a score threshold of 25.91 as identified in (25). Comparing the predicted interactions obtained by each tool in the protein coding transcriptome to SNORD126 knockdown shows that a higher proportion of the target genes predicted by snoscan are differentially expressed genes, but snoscan predicts none of the alternative splicing events upon SNORD126 knockdown (Figure 5I). SnoGloBe’s predicted interactions have the highest proportion of alternative splicing events and the second highest proportion differentially expressed genes following SNORD126 knockdown. PLEXY performs similarly to snoGloBe on the prediction of differential expression targets, but the overlap between snoGloBe and PLEXY predicted targets is small, hinting that both tools use different information and complement each other, whereas all the differentially expressed genes predicted by snoscan were also predicted by PLEXY (Figure S19). RIsearch2 has the highest number of false positives, underlining the importance of using snoRNA specific predictor to delineate high confidence interactions. Even though snoGloBe provides an enhancement and complement in the prediction of functional targets compared to other tools, not all predicted targets are affected by SNORD126 knockdown and several non-predicted targets were affected reflecting the complexity of the biological system that could vary depending on the cell line, growth conditions and propensity to secondary effects. The power of snoGloBe in predicting functionally relevant targets is evident when compared to HTRRI. In contrast to snoGloBe predictions of SNORD126 for which a total of 72 were detected as either stability and/or splicing targets following SNORD126 knockdown, only two SNORD126 interactions were identified in the HTRRI datasets, neither of which is detected as affected by the SNORD126 knockdown, emphasizing the usefulness of snoGloBe in addition to high-throughput methodologies. We conclude that snoGloBe is an efficient tool for the prediction of biologically relevant snoRNA-RNA interactions.

### SnoGloBe predicts interactions between human snoRNAs and the SARS-CoV-2 transcriptome

As another example of the utility of snoGloBe, we applied it to the SARS-CoV-2 transcriptome. It has been shown that the SARS-CoV-2 genome is heavily 2’-O-ribose methylated and interacts strongly with snoRNAs using the high-throughput structure probing methodology SPLASH (22, 60). Thus as another proof of concept for the utility and capacity of snoGloBe, we predicted the interactions between human snoRNAs and the SARS-CoV-2 genome using snoGloBe. We detected 8818 interactions between 312 snoRNAs and the SARS-CoV-2 genome, the distribution of which is shown in Figure 6A. One of the strongest interaction partners between SARS-CoV-2 and host transcripts detected experimentally using SPLASH is with the box C/D snoRNA SNORD27 (60). Although, the SPLASH experiments were carried out in Vero-E6 cells from African green monkey kidney, snoGloBe detects the SARS-CoV-2 interaction with human SNORD27 (Figure 6B), suggesting that the interaction is also relevant in human.

**Figure 6.**
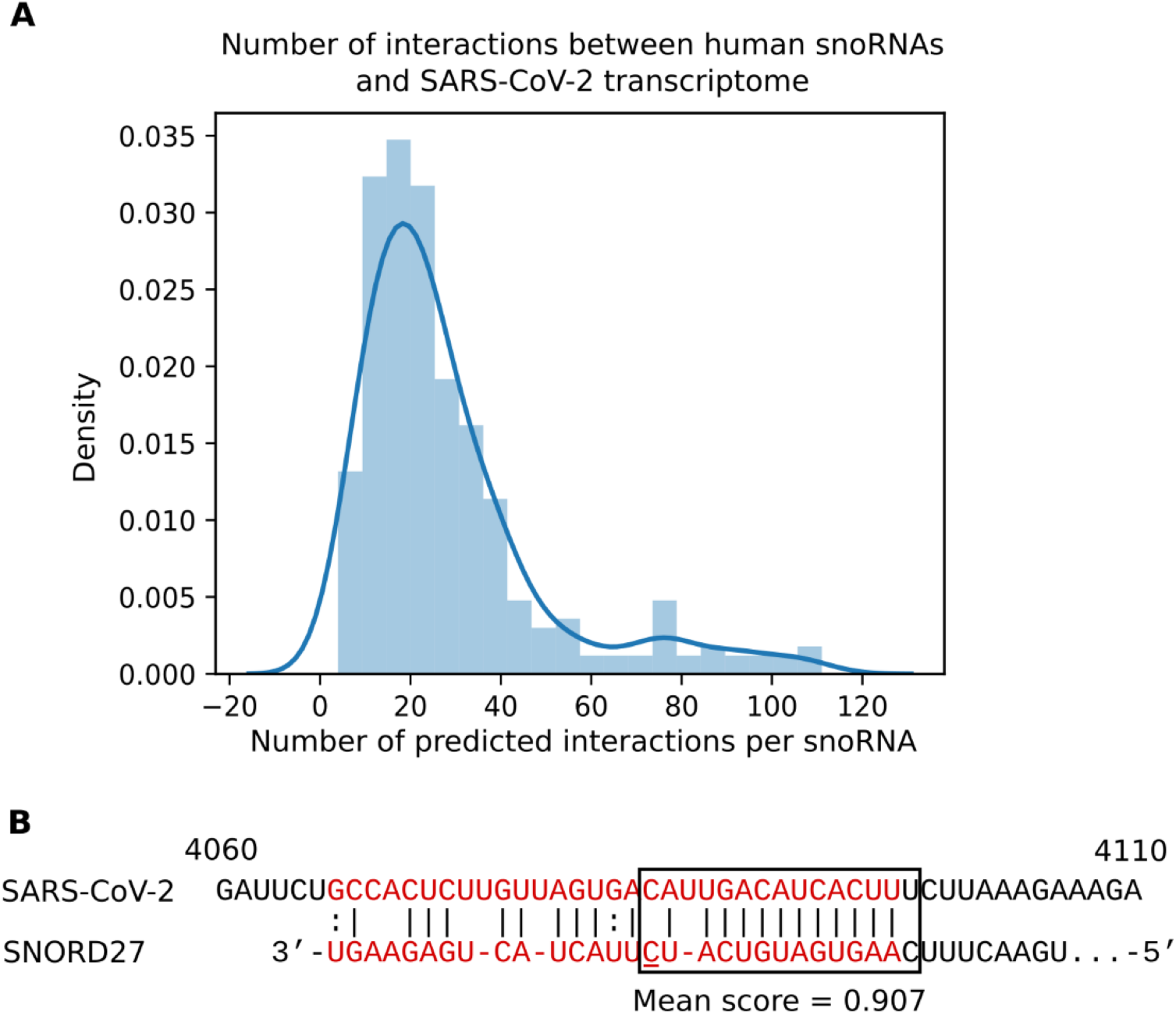
Human box C/D snoRNAs are predicted to target the SARS-CoV-2 transcriptome. A) Distribution of the number of predicted interactions per snoRNA across the SARS-CoV-2 transcriptome. Human box C/D snoRNAs have a median of 22 predicted interactions having at least 3 consecutive windows with a score greater or equal to 0.85 with SARS-CoV-2 transcriptome. B) The validated interaction between African green monkey SNORD27 and SARS-CoV-2 is also predicted with human SNORD27. The validated interaction is shown in red, the nucleotide that differs between the African green monkey and human SNORD27 is underlined. The predicted interaction is outlined by the box.

### SnoGloBe availability and usage

The snoGloBe code is written in Python using the machine learning package scikit-learn (37). It is freely available and can be downloaded from https://github.com/scottgroup/snoGloBe. Users must provide a file with the sequences of the snoRNAs of interest, the sequences of whole chromosomes, an annotation file in gtf format and a file with the potential target identifiers to scan for snoRNA interactions. Detailed instructions are available in the help manual.

## DISCUSSION

Motivated by the continually increasing number of examples of snoRNAs interacting with noncanonical targets using diverse regions within but also without the ASE (Figure 1) as well as by the diversity in RNAs targeted by snoRNAs (Figure 2A), we built snoGloBe, a box C/D snoRNA interaction predictor that considers the whole snoRNA and any type of RNA target. SnoGloBe is a gradient boosting classifier that takes into account the sequence of the snoRNA and of its potential target as well as the position in the snoRNA and the type and position in the potential target. Compared to general use RNA-RNA interaction predictors that consider only sequence complementarity and the interaction stability of the duplex, snoGloBe performs considerably better, suggesting that considering snoRNA and target features enhances the prediction. SnoGloBe also performs better than the snoRNA-specific predictors PLEXY and snoscan which were not built to predict interactions outside of the snoRNA ASEs. Many such non-ASE interactions are detected in HTRRI datasets and some have been extensively validated for individual snoRNAs (Figure 1B), limiting the scope of the PLEXY and snoscan predictors, even though snoscan performs quite well on some known noncanonical interactions. Interestingly a subset of positive examples (32%) is not found by snoGloBe and while this proportion is considerably lower than for all other predictors considered (they miss >65% of positives in the test set, Figure 3C), there is still room for improvement. This subset involves mostly HTRRI and interactions displaying bulges in the base pairing in one or both members of the interaction, which are more difficult to accurately identify and will require different approaches and likely larger training datasets for machine learning approaches to accurately predict them.

The study of the snoGloBe predicted interactions in human is very interesting and opens numerous research avenues that will likely lead to important insights into snoRNA function. Dozens of snoRNAs display profiles supporting the non-uniform distribution of predicted targets in pre-mRNA, with hundreds or even thousands of targets enriched in common regulatory elements such as PPTs, 5’ or 3’ splice sites and 5’ or 3’ UTRs (Figures 4, 5, S13, S14, S16). Each such snoRNA target profile will require in depth integrative analysis to consider the possible functionality, molecular mechanism and ultimately cellular outcome of the collective regulation of these targets by the snoRNA. We began such studies for SNORD50B, SNORD22 and SNORD126, all three of which display strong enrichment for binding to specific regulatory elements, respectively 5’ UTRs, PPTs/3’ splice sites and 3’ of introns. Manual review of their predicted targets led to the identification of a subset of such binding events overlapping alternatively regulated events (for example alternative 5’ UTRs for SNORD50B and alternatively spliced exons for SNORD22, Figures S15, S17). Gene ontology analysis of the targets show strong enrichment for specific biological processes and provide convincing subsets of targets to focus on. Finally the strong overlap between snoRNA predicted binding sites and ENCODE-detected RBP binding sites is important evidence of the functional relevance of the interactions, particularly as the function of the RBPs with the strongest overlap strongly supports the type of regulation likely carried out by the snoRNA. Several molecular mechanisms could explain the binding overlap of snoRNA and RBP at the same position on the same target pre-mRNA. The snoRNA could be guiding the RBP to its target as snoRNAs do for core snoRNA binding proteins such as FBL, and as has been reported for nuclear exosome components as shown in (13). However, since RNA binding motifs are known for several of the RBPs considered and because snoRNAs have not been found as enriched binding partners of all these RBPs, it is likely that the snoRNAs and some of the RBPs are competing for the same binding site. Further studies will be required to define the snoRNA-RBP relationship and its effect on the regulation of the targets. These overlapping snoRNA-RBP targets could be revealing novel levels of post-transcriptional regulation, the understanding of which will be important in health and disease.

Analysis of the newly predicted targets suggests that snoRNAs play an important role in regulating the splicing and stability of protein coding RNA. Indeed, the knockdown of a model snoRNA altered the stability and splicing of predicted targets even when tested in a single cell line and growth condition. Comparison with other tools showed that snoscan predicted interactions have a greater proportion of differentially expressed genes, followed by snoGloBe and PLEXY. All of snoscan predicted differentially expressed genes are also predicted by PLEXY, but the overlap between PLEXY and snoscan, and snoGloBe is low, indicating that they can be used together to identify more valid targets (Figure S19). SnoGloBe’s predicted interactions have the highest proportion of alternative splicing events (Figure 5I). SnoRNA-RNA interactions are short, which is reflected in their minimal free energy. Hence, when using general RNA-RNA interaction predictors, the minimal free energy threshold can’t be set to a very stringent value, resulting in a vast number of reported interactions, emphasizing the importance of snoRNA specific tools to help narrow the search and rapidly identify high confidence candidates. Meanwhile, by considering the ensemble of the data generated by snoGloBe, certain missed targets could be identified by virtue of independent neighbourhood analysis or similarity of their effects on cellular phenotype. Clearly most targets need to be ultimately validated experimentally and datasets like HTRRI will remain valuable resources. However, the limitation of most experimental approaches including the consideration of too few cell types and growth conditions as well as the cost and time involved makes them less useful for uncovering new targets and condition-specific binding events. Additionally, both the HTRRI datasets and snoGloBe likely predict interactions that could occur in the cell but have no functional consequences. SnoGloBe will continue to be improved in an iterative cycle as more functionally validated snoRNA-RNA interactions are identified.

Overall, while HTRRI datasets have collectively been generated in a handful of cell lines, considerable time and money would be required to explore normal human tissues and diverse conditions using these methodologies. SnoGloBe predictions will be instrumental in filling the gap by providing rapid predictions for snoRNA interactions that can then be further investigated to better understand cellular functionality. Our additional demonstration that snoGloBe can be used to investigate the interactions between snoRNAs and viral transcripts (Figure 6), further widens its scope and utility.

### DATA AVAILABILITY

All raw and processed sequencing data generated in this study have been submitted to the NCBI Gene Expression Omnibus (GEO (41); https://www.ncbi.nlm.nih.gov/geo/) under accession number GSE184173.

## ACKNOWLEDGEMENTS

The authors would like to thank members of the Scott and Abou-Elela groups for helpful discussions and Compute Canada for providing state-of-the-art computing infrastructures. SAE and MSS are members of the Centre de Recherche du Centre Hospitalier de l’Université de Sherbrooke.

## FUNDING

This work was supported by a Canadian Institutes of Health Research grant [PJT 153171 to MSS and SAE] and a team FRQ-NT grant to MSS and SAE. GDF was supported by a NSERC Doctoral scholarship. MSS holds a Fonds de Recherche du Québec – Santé (FRQ-S) Research Scholar Senior Career Award.

## CONFLICT OF INTEREST

The authors declare no competing interests.

---

**Figure S1:**
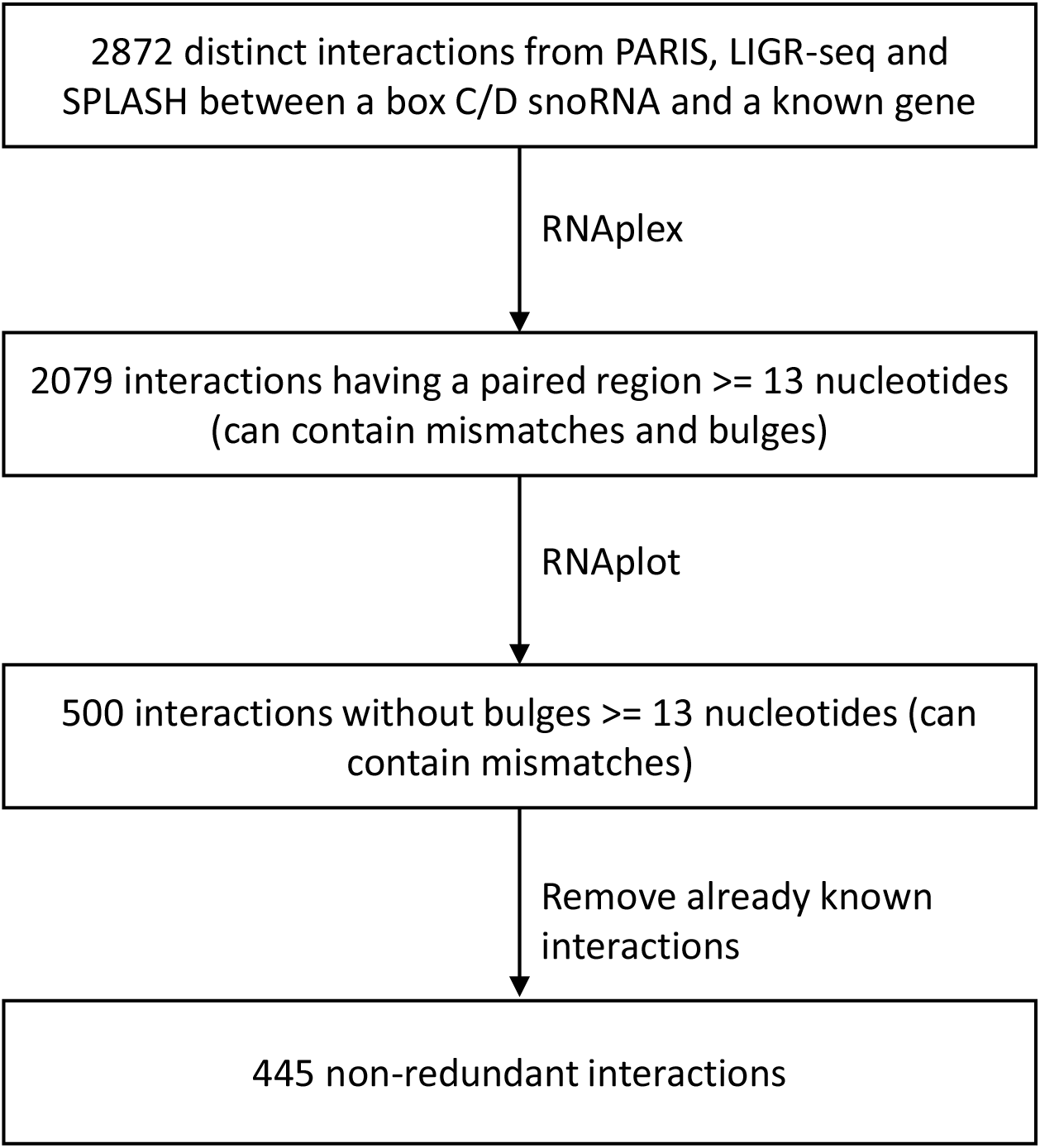
Overview of the filtering procedure starting with all distinct interactions detected in at least one HTRRI dataset involving a snoRNA to obtain the final HTRRI formatted datasets. The number of interactions remaining at each step is indicated.

**Figure S2:**
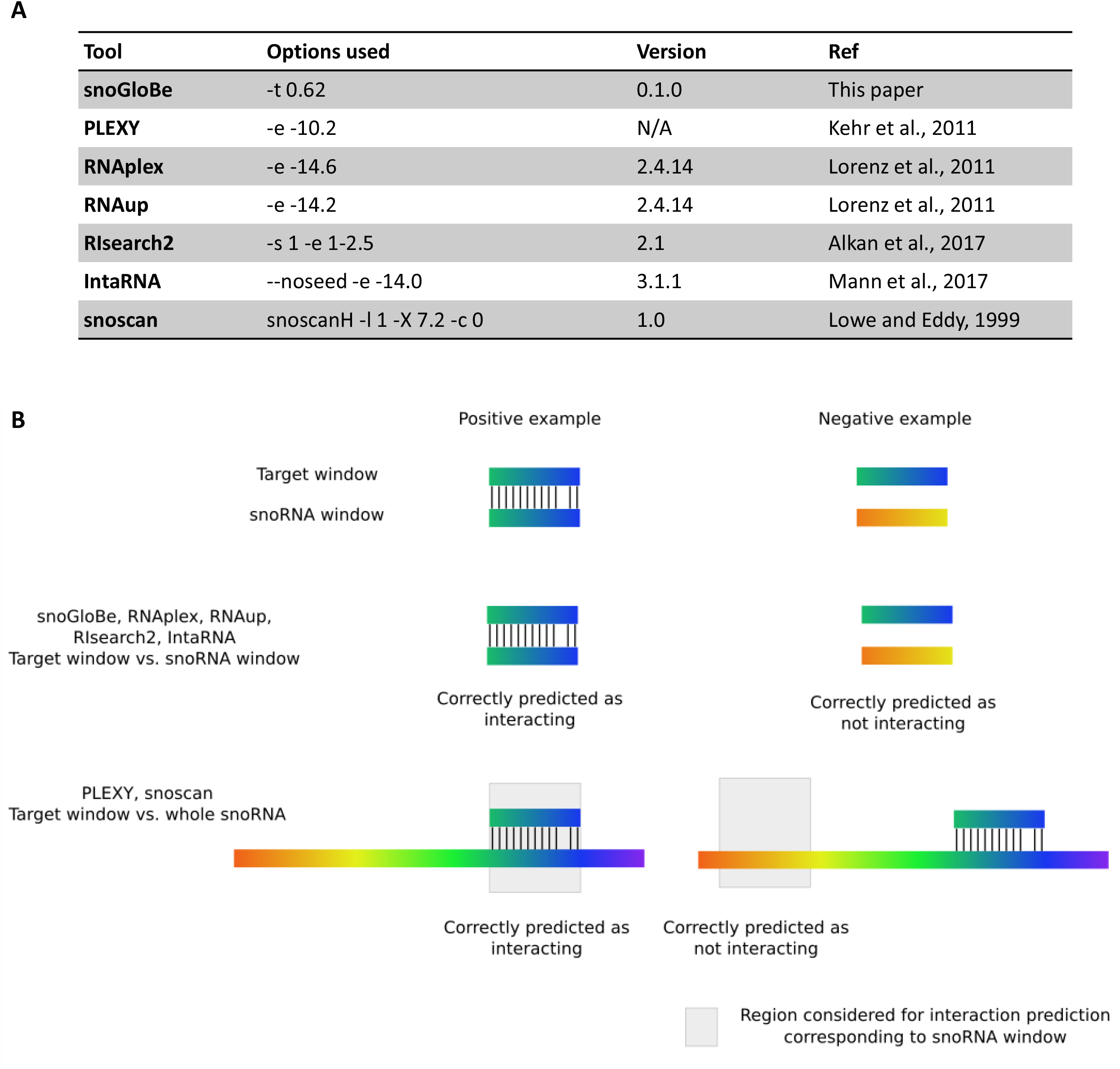
(A) List of tools compared for the prediction of C/D snoRNA-RNA interactions as well as their parameter values used, version and paper reference. (B) Strategy used to determine the prediction status of snoRNA-target window pairs for each tool. PLEXY and snoscan require the entire snoRNA, so only the interactions predicted in the snoRNA window region were considered to ensure fairness between tools.

**Figure S3:**
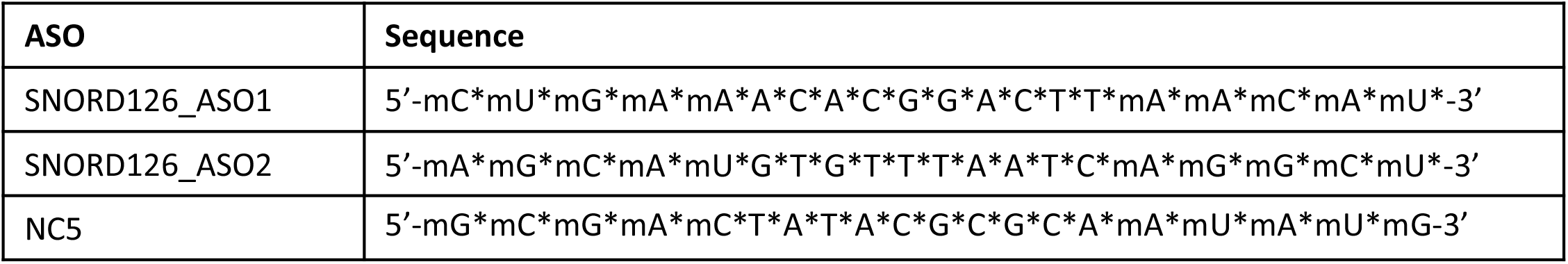
List of ASO sequences used for SNORD126 knockdown and negative control (NC5). * means phosphorothioate backbone, m means 2’-O-methoxyethyl.

**Figure S4:**
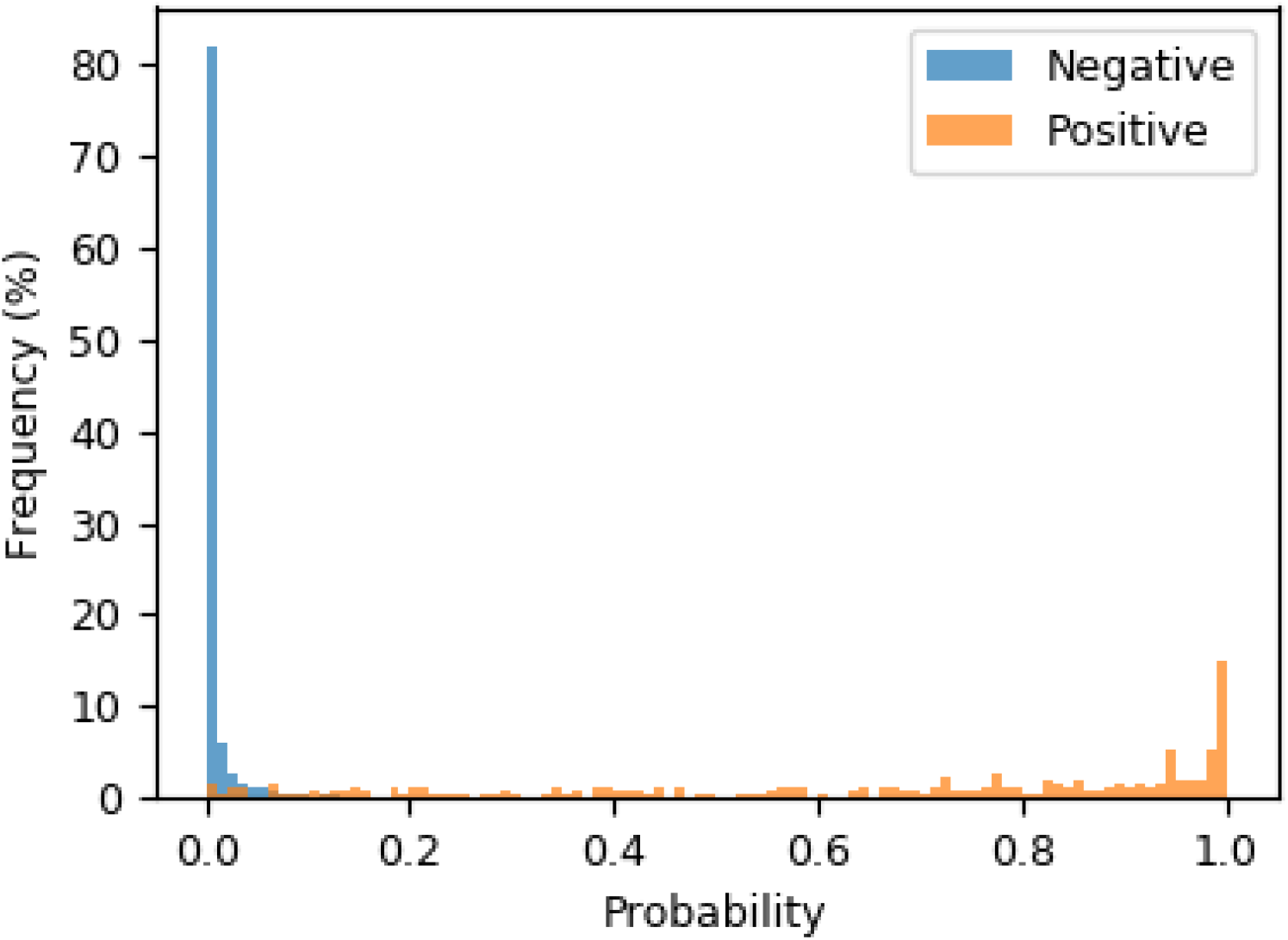
Distribution of the scores output by snoGloBe for the positive and negative examples in the test set.

**Figure S5:**
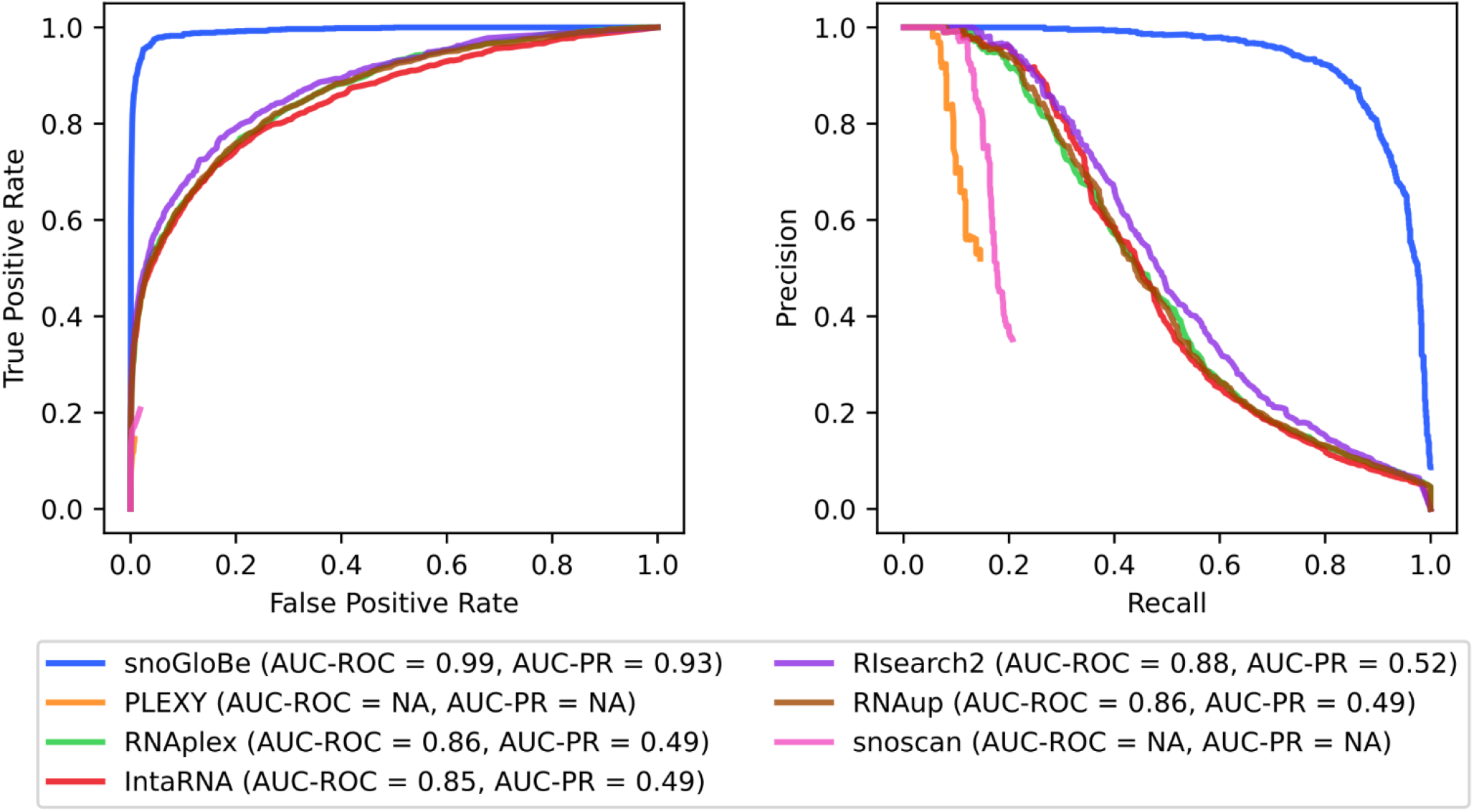
Receiver Operator Characteristic (ROC) and (B) Precision-Recall (PR) curves of different tools calculated on the training set. The corresponding area under the curves (AUC) are indicated in the legend.

**Figure S6:**
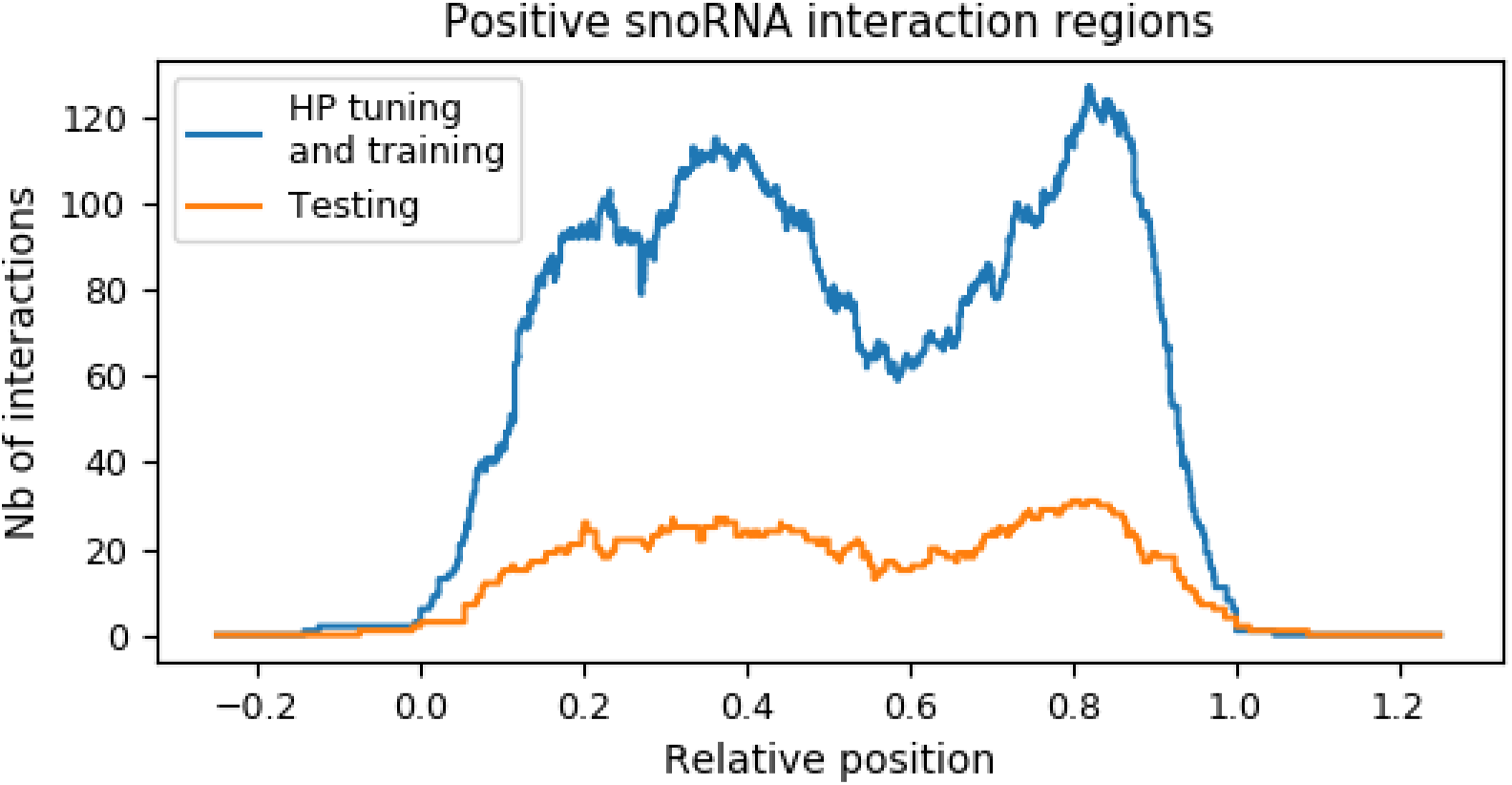
Profiles representing the number of predicted interactions from the hyperparameter tuning and training sets (blue) or the test set (orange) that involve specific positions in the snoRNA, for all snoRNAs considered simultaneously.

**Figure S7:**
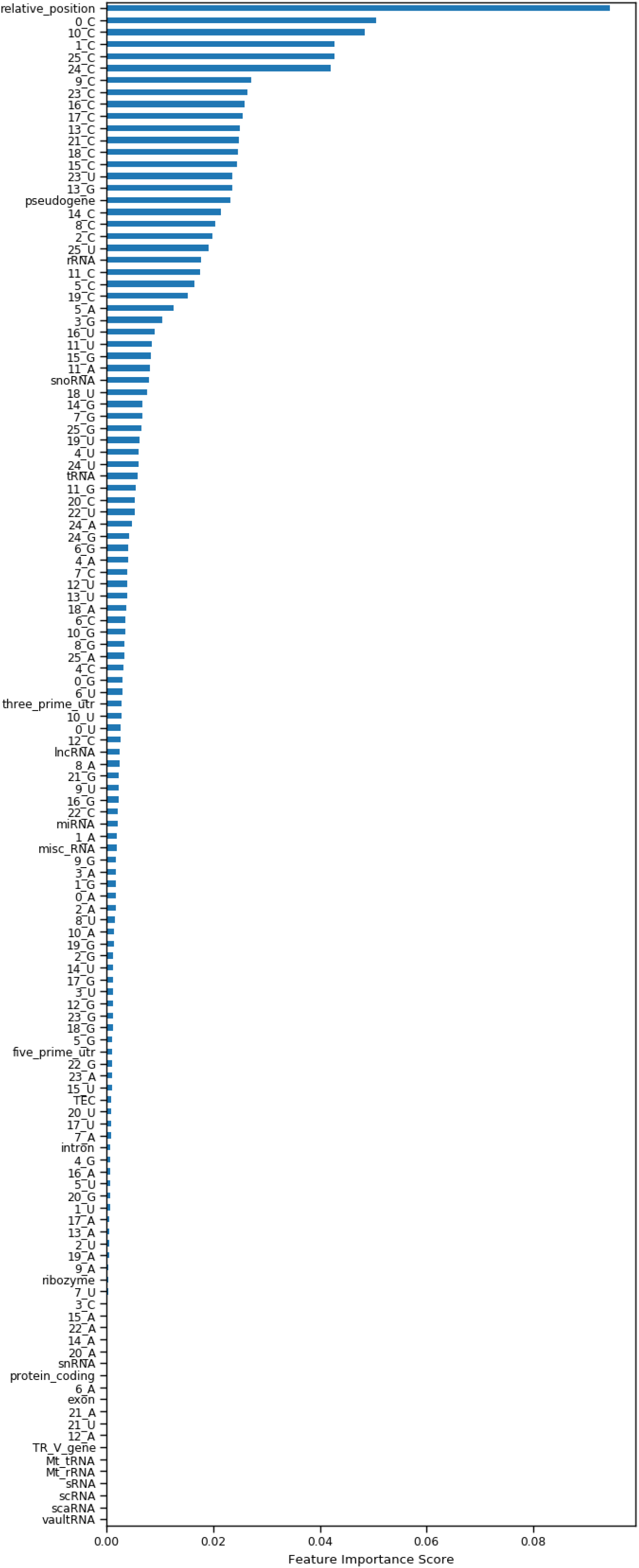
Score measuring the importance of each input feature. A higher score means that the feature has more importance in the overall classification of a potential interaction. The features identified by a number followed by a nucleotide are the importance on each nucleotide at a given position of the window. The nucleotides 0-12 represent the snoRNA window and 13-25 the target window, both in a 5’-3’ manner.

**Figure S8:**
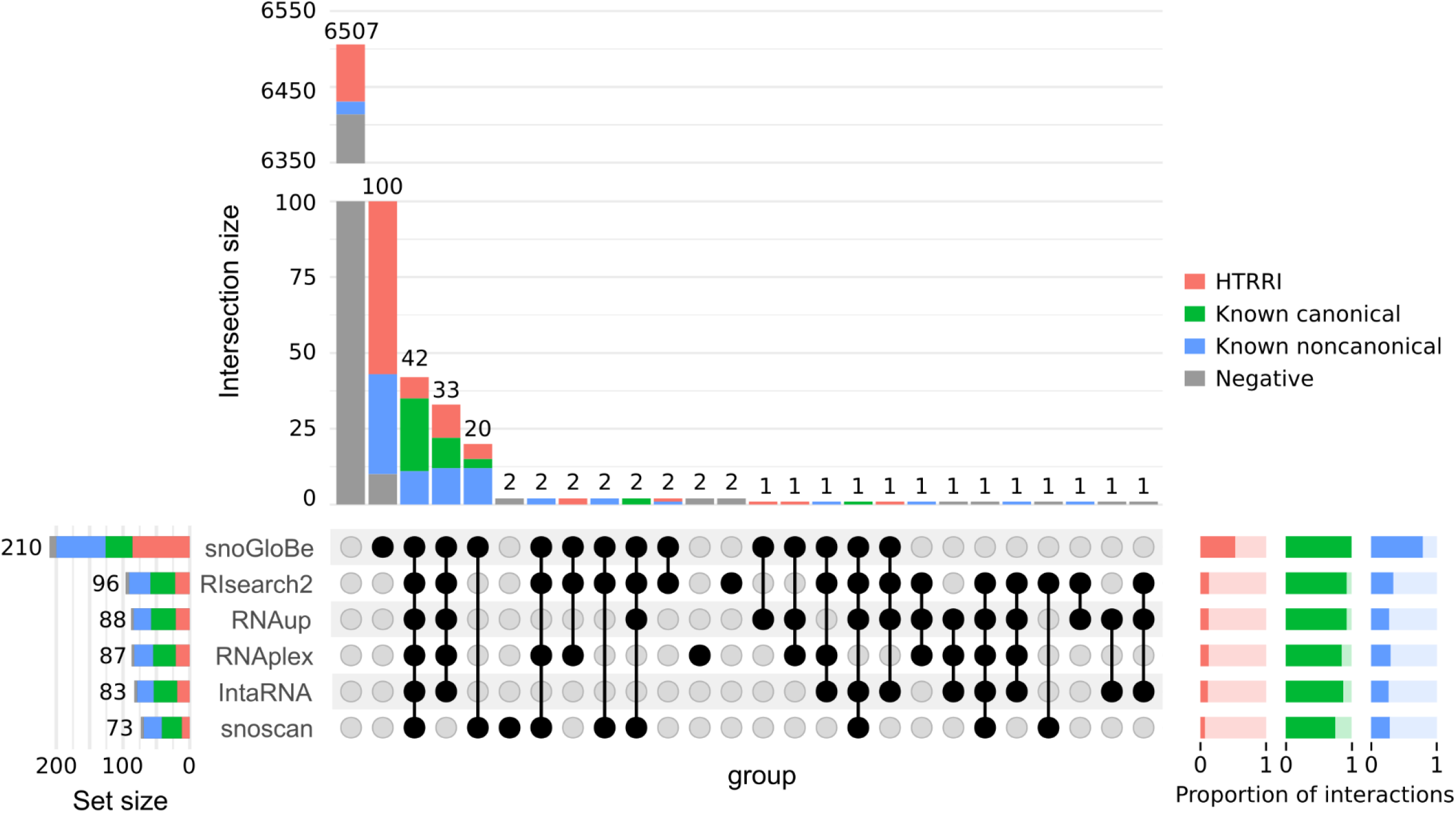
Upset plot indicating the number of predicted interactions from each category for all predictors compared considering all the box C/D snoRNAs. The color legend on the right indicates the different interaction types. The proportion of positive examples from each category predicted as positive (dark color) and negative (light color) by each tool is shown in the bottom right. This figure differs from Figure 3D by showing the interactions from all the snoRNAs from the test set, whereas the Figure 3D only shows the snoRNAs that have non-degenerated boxes D and D’, which are the snoRNAs that we used with PLEXY. Here, PLEXY is not shown.

**Figure S9:**
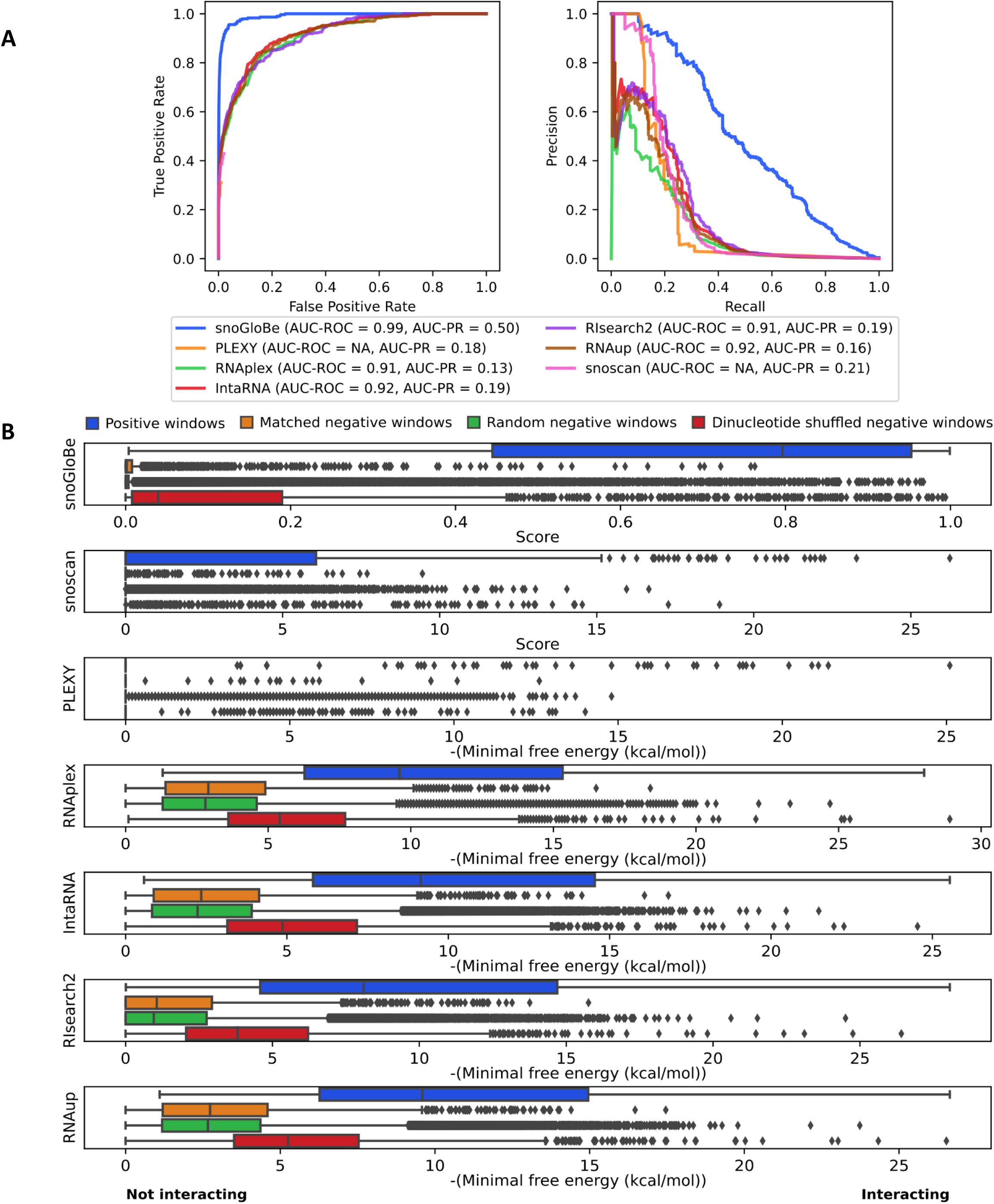
(A) Receiver Operator Characteristic (ROC) and Precision-Recall (PR) curves of different tools calculated on the test set to which were added negative examples to get 1000 random negative examples and 7 negative examples obtained by dinucleotide shuffling for each positive example. The corresponding area under the curves (AUC) are indicated in the legend. (B) Distribution of the score or minimal free energy obtained by every example for each tool divided by category in the following order: positive windows (blue), matched negative windows (orange), random negative windows (green) and dinucleotide shuffled negative windows (red). The scale of the minimal free energy is reversed to get the windows predicted as not interacting on the left and those predicted as interacting on the right for all tools.

**Figure S10:**
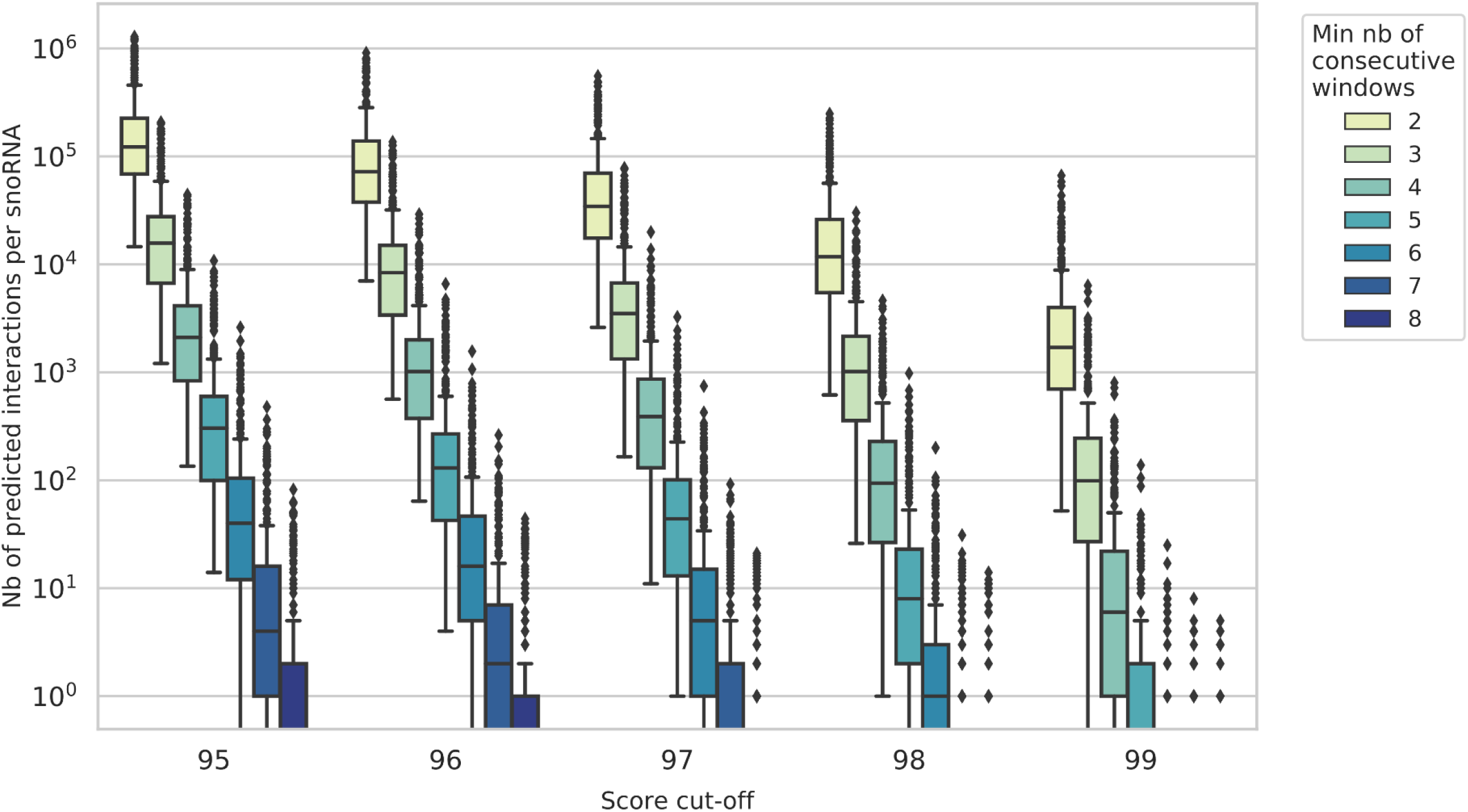
Boxplots depicting the distributions of the number of predicted interactions as a function of the parameter values used. The parameters considered are the number of predicted interactions per snoRNA (y axis), the score cut-off (x axis) and the minimum number of consecutive windows (different colors, with legend on the right).

**Figure S11:**
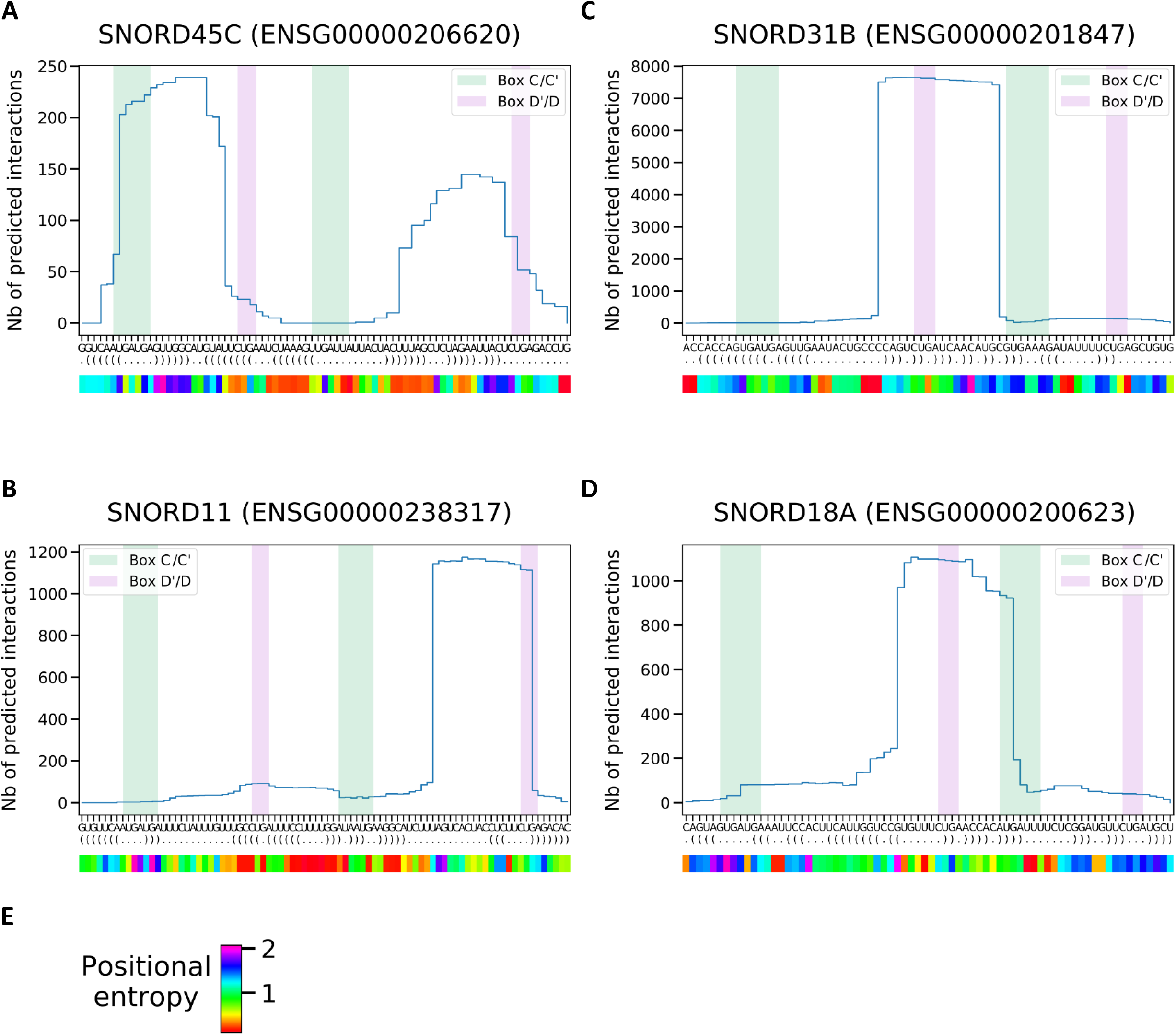
Profiles measuring the number of predicted interactions involving each position of the snoRNAs SNORD45C (A), SNORD11 (B), SNORD31B (C) and SNORD18A (D). The green and pink highlighting in the profiles represent respectively the positions of the C’/C and D’/D boxes. The predicted folding of each snoRNA, as predicted by RNAplot, is shown using the dot-bracket format. The panel below each profile represents the positional entropy predicted by replot from the ViennaRNA package for each position of the snoRNA. The color legend for the position entropy score is given in E.

**Figure S12:**
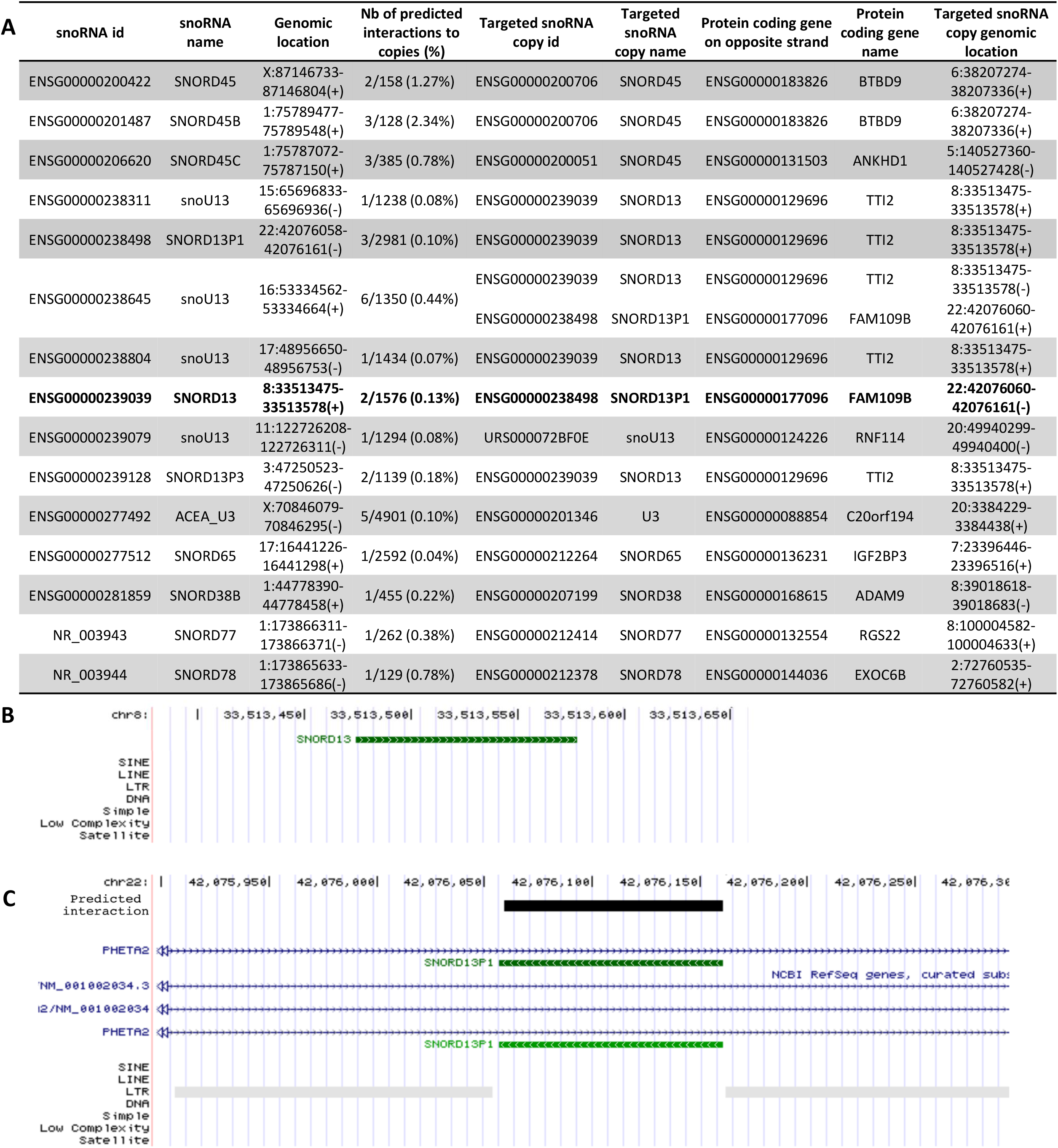
(A) Table describing the targets that are full-length copies of snoRNAs. We found that 15/312 snoRNAs considered have full-length targets antisense to a protein coding gene, likely representing transposition or recombination events copying the snoRNA to another genomic location. The identifiers and genomic location of the snoRNA and the targeted copies are given in the table as well as the identifiers of the protein coding genes. The column ‘Nb of predicted interactions to copies’ represents the number of interactions to full-length copies out of all interactions to all targets of the snoRNA. (B-C) Screenshots of the UCSC genome browser of a snoRNA (B) and copied snoRNA on the opposite strand of the protein coding gene PHETA2 (C). The copied snoRNA is surrounded by LTR regions. The snoRNA and the copy are located on different chromosomes. The predicted interaction on the copied snoRNA is shown in black.

**Figure S13:**
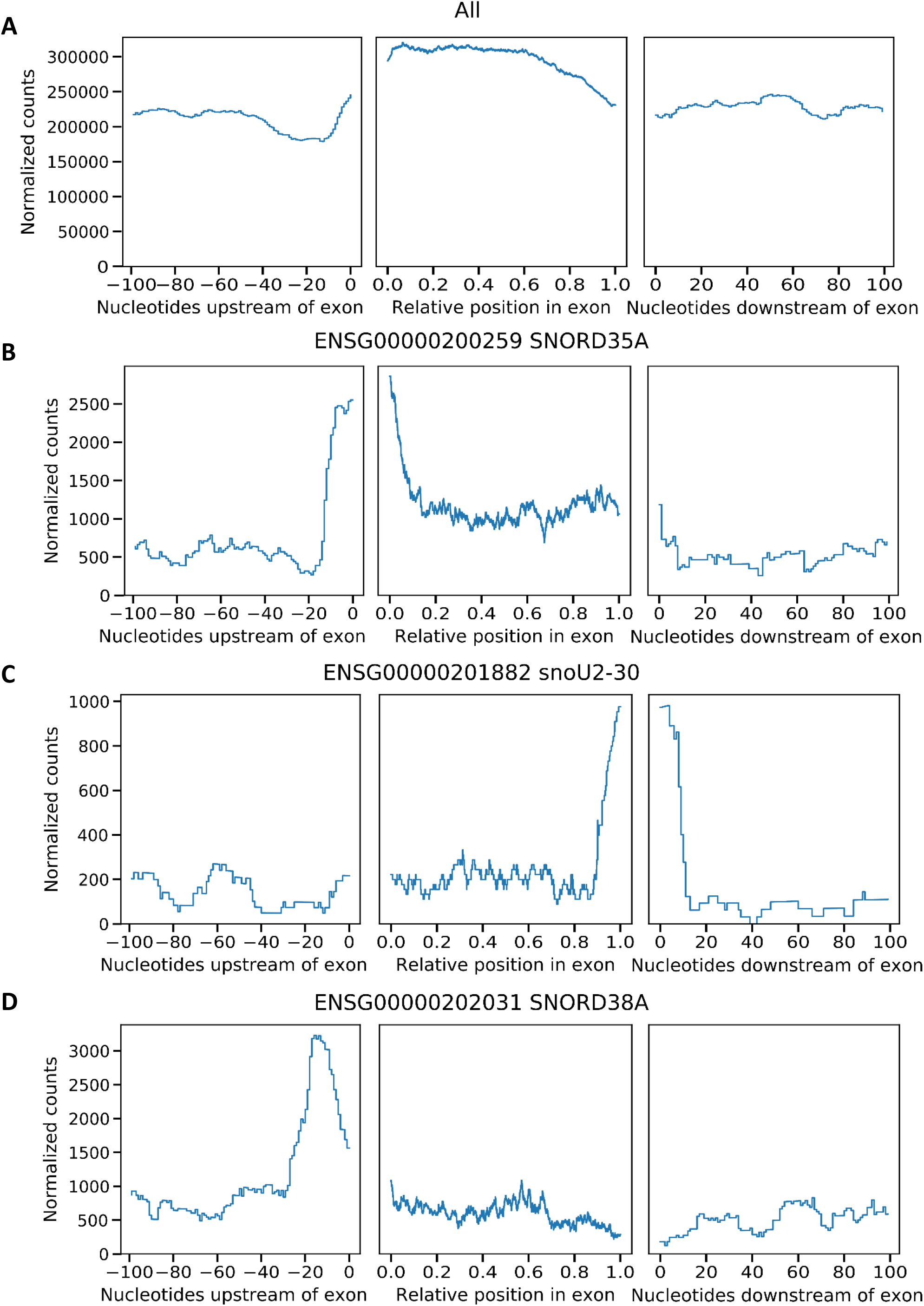
Profiles measuring the number of predicted interactions as a function of position with respect to the termini of introns and exons for all targets of all snoRNAs (A), SNORD35A (B), snoU2-30 (C) and SNORD38A (D).

**Figure S14.**
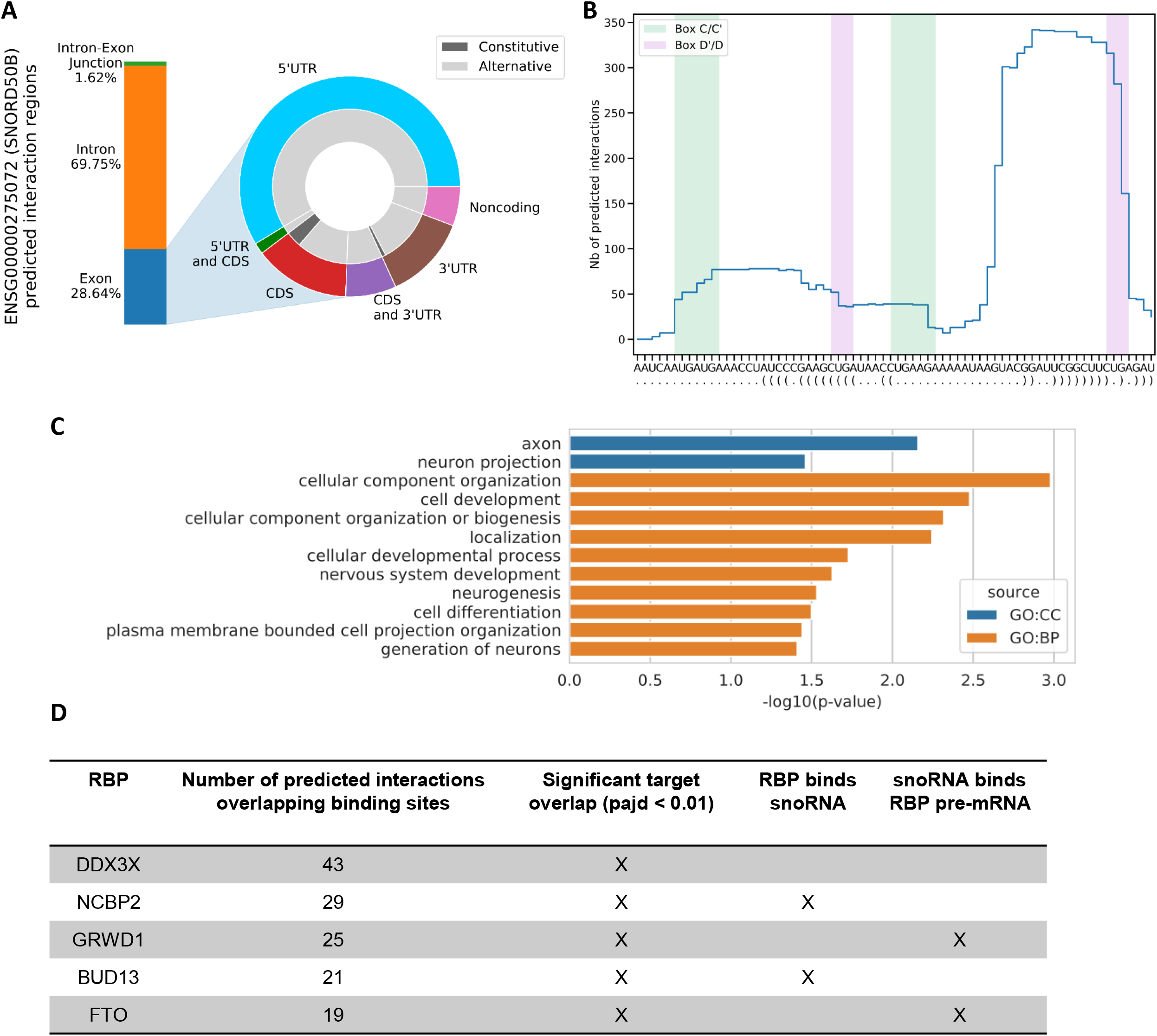
SNORD50B displays a positional enrichment of binding on functionally related targets which are targeted by related RBPs. (A) SNORD50B targets are enriched in exons and more specifically in 5’ UTRs compared to the transcriptome distribution (Fig. 4 C and E) as shown using a stacked bar plot showing the distribution of its targets across introns and exons (left) with a doughnut plot showing the breakdown for different exonic elements (right). (B) SNORD50B binds its targets using mainly its box D ASE, as shown with a positional profile measuring the number of targets each position of the snoRNA is predicted to bind. (C) SNORD50B predicted targets are enriched in specific gene ontology terms (CC: cellular component, BP: biological process, MF: molecular function) indicated using a horizontal bargraph. (D) SNORD50B displays strong association with specific RBPs. Table showing the number of predicted SNORD50B interactions that overlap binding sites for the indicated RBPs. (All 5 RBPs indicated have a significant overlap). In addition, 2 of the RBPs bind SNORD50B according to eCLIP experiments and SNORD50B is predicted to bind the pre-mRNA of 2 of the RBPs.

**Figure S15.**
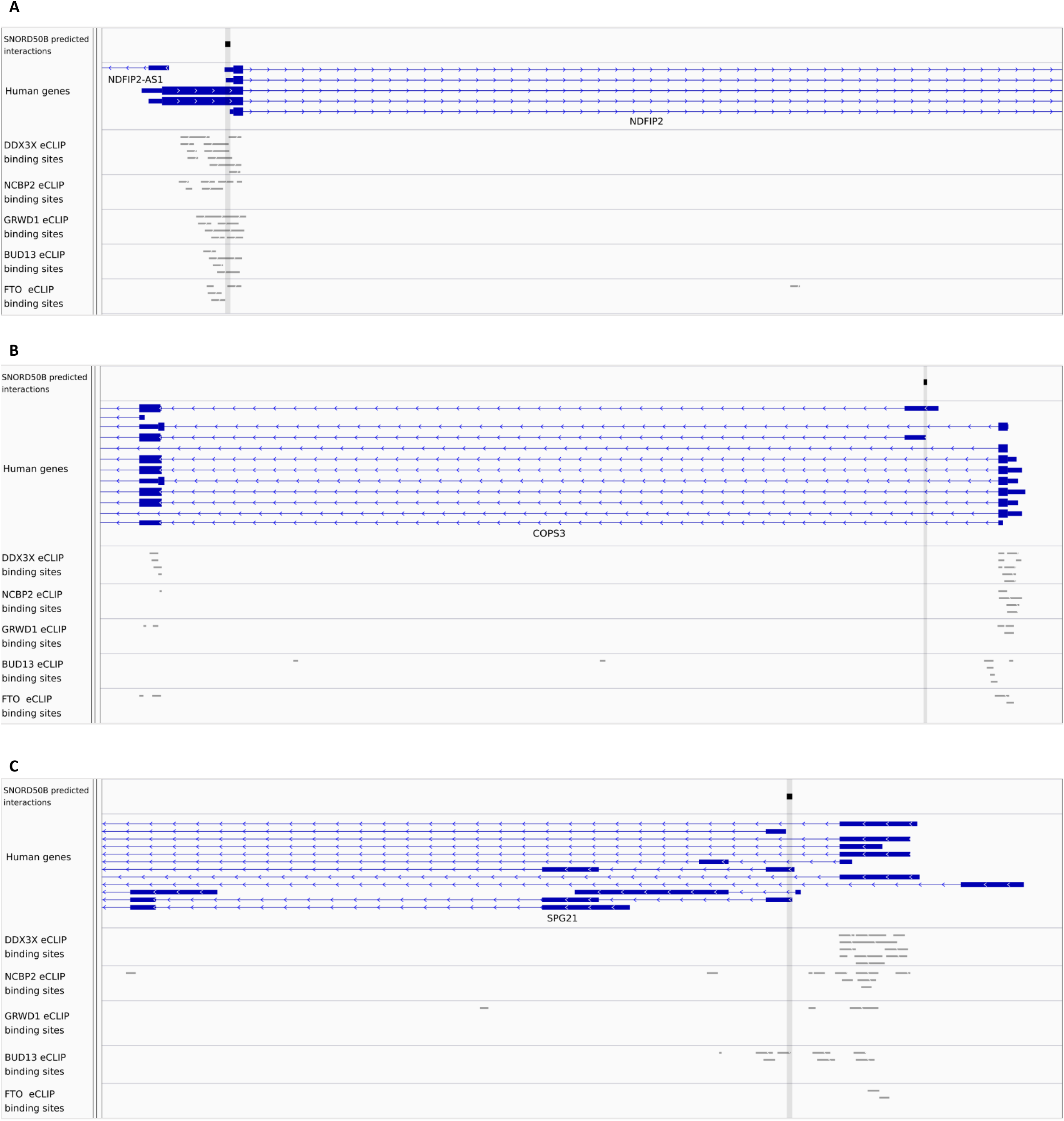
Examples of SNORD50B binding sites on alternative 5’ UTRs. SNORD50B binding sites are enriched in 5’ UTRs many of which are alternative including those in (A) NDIFP2, (B) COPS3 and (C) SPF21, as shown with genome browser screenshots. In each case, the top track displays the predicted binding position. The Human genes track indicates the architecture of the different isoforms encoded for these genes for the positional window chosen and the bottom tracks show the binding sites for the 5 RBPs indicated in Figure S14D according to ENCODE eCLIPs.

**Figure S16.**
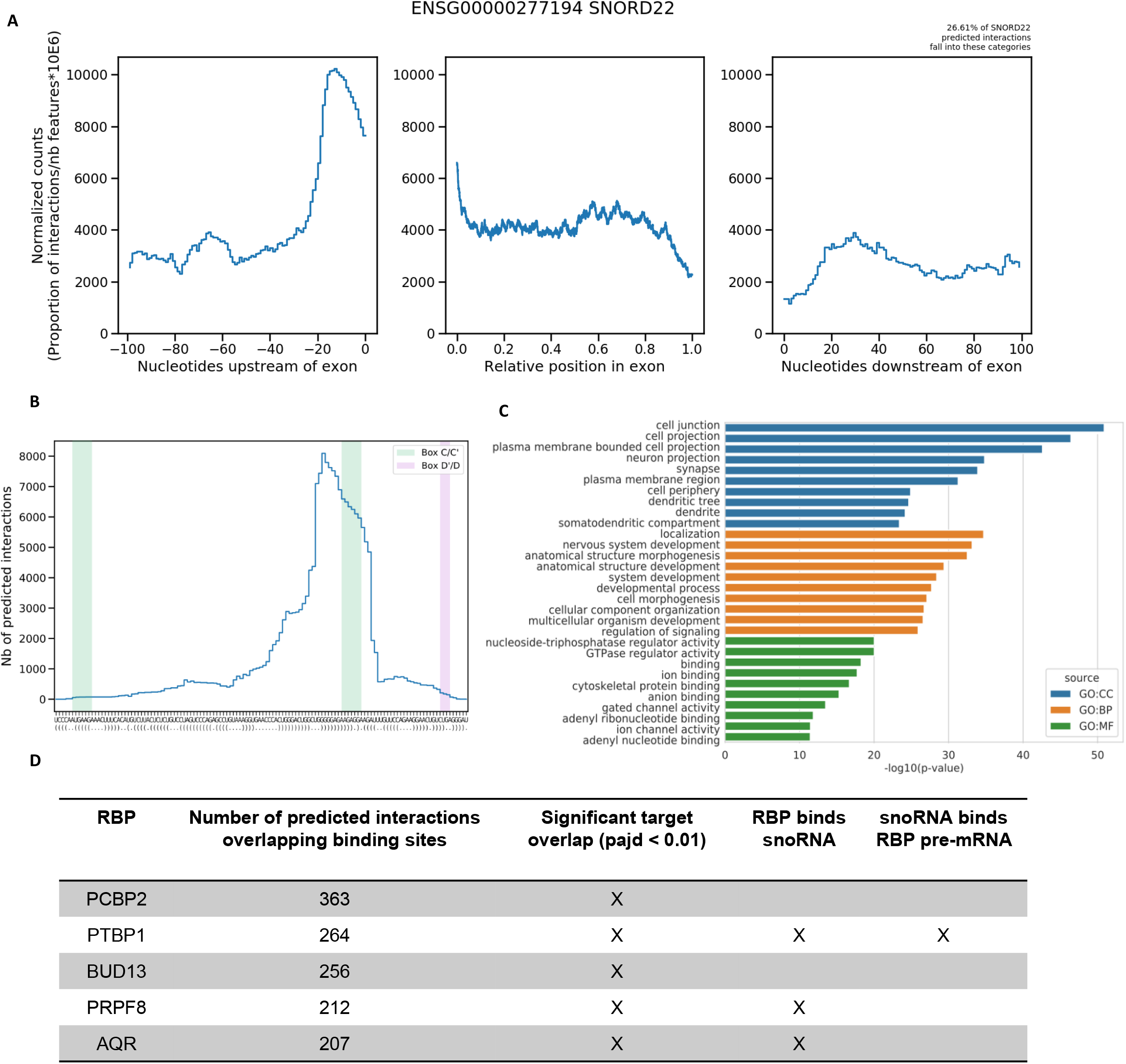
SNORD22 displays a positional enrichment of binding on functionally related targets which are targeted by related RBPs involved in splicing. (A) SNORD22 targets are enriched in 3’ SSs as shown using a positional profile covering the last 100 nt of introns, the relative position in exons and the first 100 nt in introns. (B) SNORD22 binds its targets using a region overlapping its box C’. (C) SNORD22 predicted targets are enriched in specific gene ontology terms (CC: cellular component, BP: biological process, MF: molecular function) indicated using a horizontal bargraph. (D) SNORD22 displays strong association with specific RBPs. Table showing the number of predicted SNORD22 interactions that overlap binding sites for the indicated RBPs. (All 5 RBPs indicated have a significant overlap). In addition, 3 of the RBPs bind SNORD22 according to eCLIP experiments and SNORD22 is predicted to bind the pre-mRNA of 1 of the RBPs.

**Figure S17.**
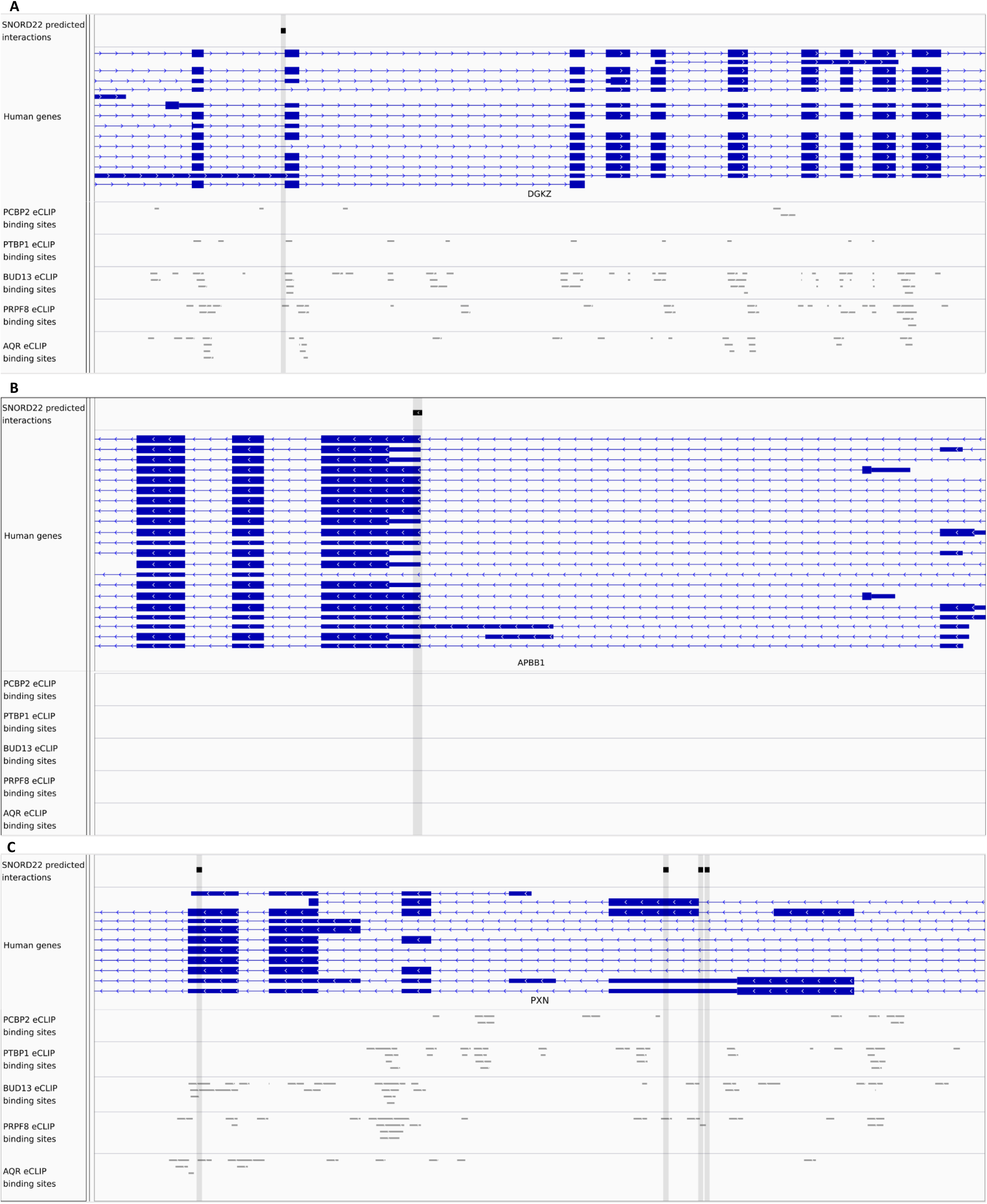
Examples of SNORD22 binding sites on alternative 3’SSs. SNORD22 binding sites are enriched in 3’ SSs many of which are alternative including those in (A) DGKZ, (B) APBB1 and (C) PXN. Screenshots are as described in Figure S15 and the RBP tracks correspond to those from Figure S16D.

**Figure S18:**
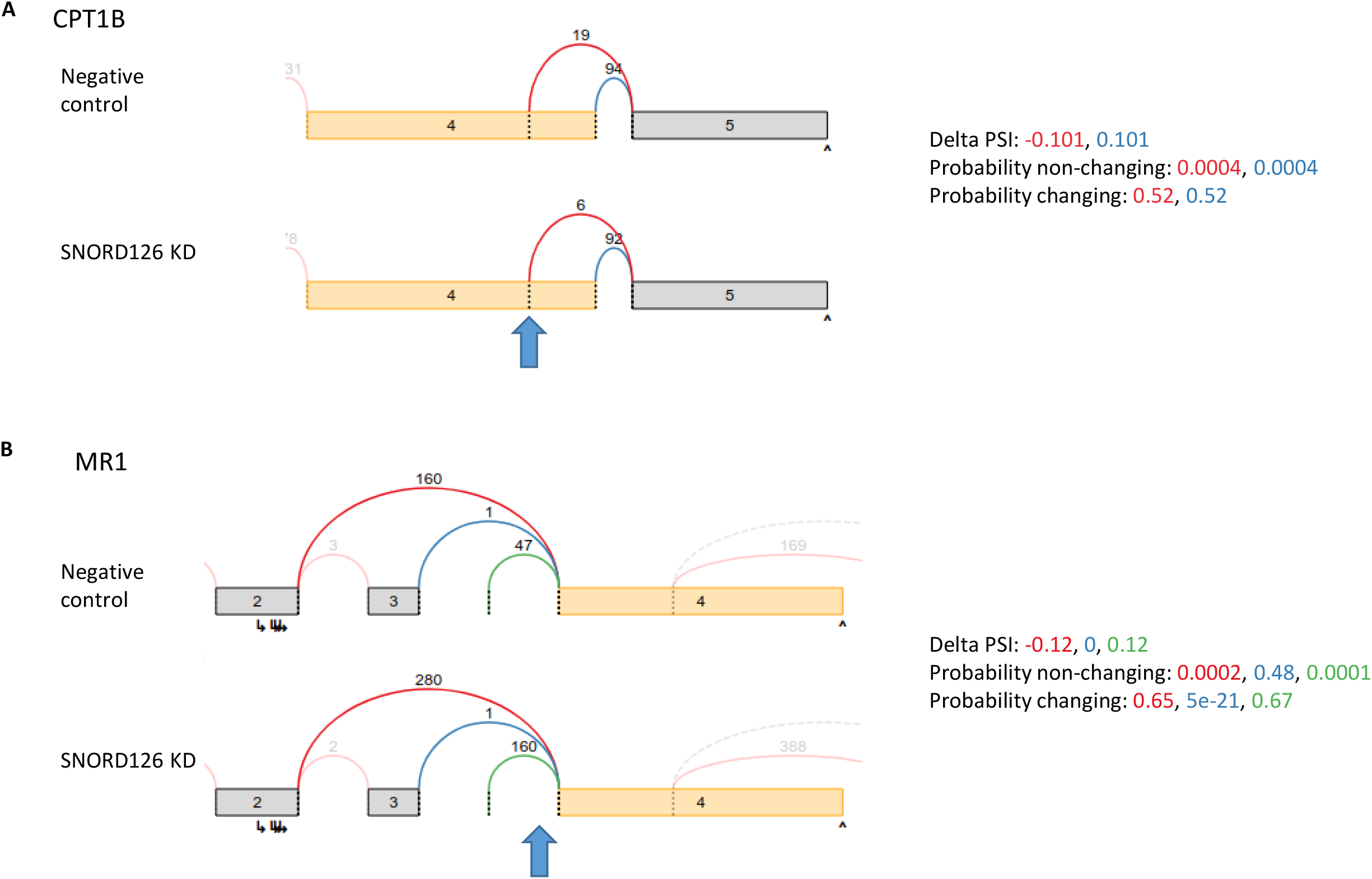
Examples of splicing events affected by the knockdown of SNORD126. Both CPT1B (A) and MR1 (B) display differential splicing following the knockdown (KD) of SNORD126 as shown using sashimi plots. The blue arrows represent the predicted interaction region. The colored arcs represent different splice junctions with the number of reads supporting them. Statistics of the splicing event are given on the right.

**Figure S19:**
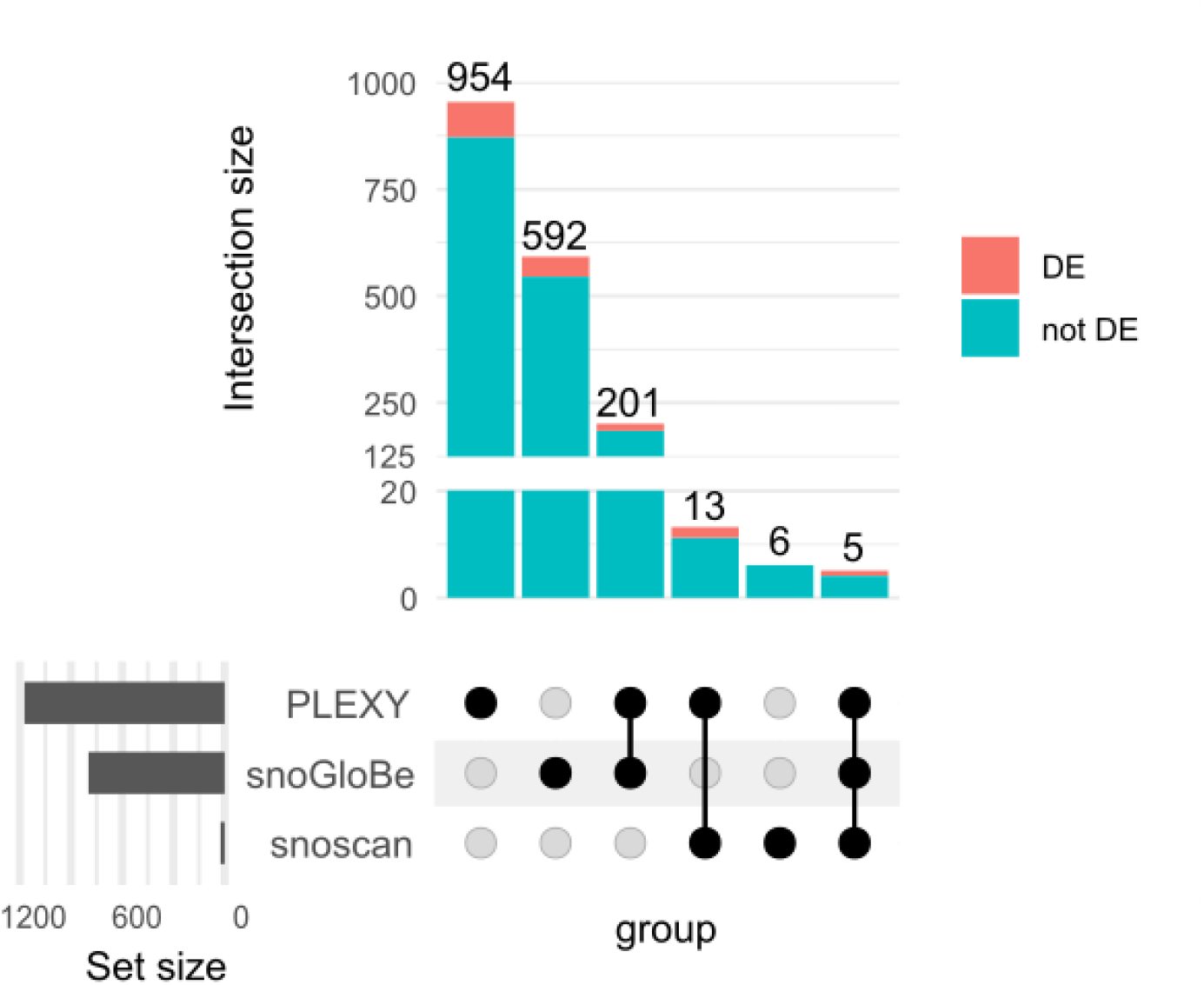
Overlap between the genes having at least one interaction predicted by PLEXY, snoGloBe and/or snoscan and whether they are differentially expressed (DE) or not (not DE).

## Notes

### Competing Interest Statement

The authors have declared no competing interest.

